# Conformational Flexibility in Neutralization of SARS-CoV-2 by Naturally Elicited Anti-SARS-CoV-2 Antibodies

**DOI:** 10.1101/2022.04.03.486854

**Authors:** Ruofan Li, Michael Mor, Bingting Ma, Alex E. Clark, Joel Alter, Michal Werbner, Jamie Casey Lee, Sandra L. Leibel, Aaron F. Carlin, Moshe Dessau, Meital Gal-Tanamy, Ben A. Croker, Ye Xiang, Natalia T. Freund

**Affiliations:** Beijing Advanced Innovation Center for Structural Biology, Beijing Frontier Research Center for Biological Structure, Center for Infectious Disease Research, School of Medicine, Tsinghua University, Beijing, China; Department for Microbiology and Clinical Immunology, Faculty of Medicine, Tel Aviv University, Israel; Department of Medicine, University of California San Diego, La Jolla, CA, USA; The Laboratory of Structural Biology of Infectious Diseases, Azrieli Faculty of Medicine, Bar Ilan University, Israel; Molecular Virology Lab, Azrieli Faculty of Medicine, Bar Ilan University, Israel; Department of Pediatrics, School of Medicine, UC San Diego, La Jolla, CA, USA; Sanford Burnham Prebys Medical Discovery Institute, La Jolla, CA, USA; Sanford Consortium for Regenerative Medicine, La Jolla, CA, USA

## Abstract

As new variants of SARS-CoV-2 continue to emerge, it is important to assess the neutralizing capabilities of naturally elicited antibodies against SARS-CoV-2. In the present study, we evaluated the activity of nine anti-SARS-CoV-2 monoclonal antibodies (mAbs), previously isolated from convalescent donors infected with the Wuhan-Hu-1 strain, against the SARS-CoV-2 variants of concern (VOC) Alpha, Beta, Gamma, Delta and Omicron. By testing an array of mutated spike receptor binding domain (RBD) proteins, cell-expressed spike proteins from VOCs, and neutralization of SARS-CoV-2 VOCs as pseudoviuses, or as the authentic viruses in culture, we show that mAbs directed against the ACE2 binding site (ACE2bs) are far more sensitive to viral evolution compared to anti-RBD non-ACE2bs mAbs, two of which kept their potency against all VOCs tested. At the second part of our study, we reveal the neutralization mechanisms at high molecular resolution of two anti-SARS-CoV-2 neutralizing mAbs by structural characterization. We solved the structures of the Delta-neutralizing ACE2bs mAb TAU-2303 with the SARS-CoV-2 spike trimer and RBD at 4.5 Å and 2.42 Å, respectively, revealing a similar mode of binding to that between the RBD and the ACE2 receptor. Furthermore, we provide five additional structures (at resolutions of 5.54 Å, 7.76 Å, 6.47 Å, 3.45 Å, and 7.32 Å) of a second antibody, non-ACE2bs mAb TAU-2212, complexed with the SARS-CoV-2 spike trimer. TAU-2212 binds an exclusively quaternary epitope, and exhibits a unique, flexible mode of neutralization that involves transitioning between five different conformations, with both arms of the antibody recruited for cross linking intra- and inter-spike RBD subunits. Our study provides new mechanistic insights about how antibodies neutralize SARS-CoV-2 and its emerging variants and provides insight about the likelihood of reinfections.

## INTRODUCTION

Within two years of the emergence of SARS Coronavirus 2 (SARS-CoV-2) in Wuhan province, the original virus strain has been completely replaced by more transmissible variants, with Omicron emerging as the latest variant of concern (VOC). In view of the unexpectedly fast rates of viral evolution, it is important to estimate the degree to which neutralizing antibodies elicited naturally following infection with the original wild type strain (Wuhan-Hu-1), are cross reactive with circulating, present and future, VOCs. This is particularly relevant considering recent reports that vaccination provides considerably less protection against SARS-CoV-2 variants than against the original strain^1-3^.

The SARS-CoV-2 antibody response has been profiled at the sequence, structural, and mechanistic level by cloning and characterizing monoclonal antibodies (mAbs) from Wuhan-Hu-1-infected convalescent donors^4-9^. However, with the emergence of variants, many of these mAbs, some of which have been approved for treatment of COVID-19 patients^10-12^, have become ineffective^13-15^, while others retain activity^16^. This indicates that some antibodies elicited by infection are more variation-sensitive than others, and that antibody breadth of specificity, and not only potency, should be considered. A finer resolution investigation of the mechanistic and functional basis of SARS-CoV-2 antibody neutralization is therefore needed to predict the effect viral modifications will have on antibody activity, and to estimate the degree of protection from SARS-CoV-2 reinfection and breakthrough infection. Moreover, studying the molecular recognition of naturally elicited neutralizing antibodies in the context of heterologous viruses can reveal sites with a lower tendency to variation.

We have previously identified a panel of neutralizing SARS-CoV-2 antibodies derived from two COVID-19 survivors who were infected in Israel in March 2020, likely with the Wuhan-Hu-1 strain^9^. Seven of these mAbs (TAU-1109, -1145, -2189, -2212, -2230, -2303, and -2310) exhibited potent SARS-CoV-2 neutralizing activity, while the activity of another two (TAU-1115, and TAU-2220) was less potent. All the mAbs, except one, bind soluble receptor binding domain (RBD) and spike with high affinity. The exception, TAU-2212, which is one of the most potent neutralizing mAbs, binds an unknown conformational surface, and binding can only be detected when the spike protein is expressed on viral particles or cells. While exhibiting neutralizing activity against the Wuhan-Hu-1 strain, the cross-reactivity of these mAbs with SARS-CoV-2 VOCs is unknown.

The present study was designed to investigate the breadth of specificity and structural basis of the neutralizing activity of our previously isolated mAbs, in the context of the emerging variants Alpha (B.1.1.7)^17^, Beta (B.1.351)^18^, Gamma (P.1 or B.1.1.28.1)^19^, Delta (B.1.617.2)^20,21^ and Omicron (B.1.1.529)^22^. The results indicate that the most potent mAbs in our panel are predominantly directed against the ACE2 binding site (ACE2bs) supersite and are also the ones most sensitive to viral diversification. To understand the basis of neutralization and escape at the atomic level, we used cryo electron microscopy (cryoEM) and X-ray crystallography to determine structures of the two mAbs: TAU-2303 and TAU-2212. The results indicate that the interaction of TAU-2303 Fab and SARS-CoV-2 RBD resembles that of the ACE2 receptor with RBD in the “up” RBD conformation. In contrast, mAb TAU-2212 exhibits an unusual recognition flexibility type of binding involving five different possible conformations, with 1-3 Fabs binding to one spike trimer, or cross-linking adjacent trimers by forming both intra- and inter-spike contacts, and favoring RBD in the “down” position. Our study provides important mechanistic and structural insights about neutralization of SARS-CoV-2 VOCs by natural antibodies, together with molecular modeling predictions of mAb interactions with the Omicron variant.

## RESULTS

### Antibodies binding at the ACE2bs are more sensitive to viral mutations

The results of our previous study indicated that mAbs TAU-1145, -2189, -2230, and - 2303 compete with ACE2, and are therefore defined as ACE2bs mAbs, while mAbs TAU-1109, -1115, -2220, -2310 do not compete with ACE2, and are therefore defined as non-ACE2bs^9^. The last neutralizing mAb, TAU-2212, does not bind soluble SARS-CoV-2 antigens (RBD or spike) by ELISA and recognizes an unknown epitope^9^. To examine the recognition of the RBDs from SARS-CoV-2 VOCs by our previously isolated mAbs, we generated soluble RBD proteins containing the mutations identified in the Alpha, Beta, Gamma, Delta and Omicron strains (Extended Data Table 1) and tested the binding to the eight mAbs that originally exhibited strong binding to the wild type strain RBD^9^. With the exception of Alpha, we observed a general reduction in binding efficiency of the ACE2bs mAbs to all the VOCs (Fig. 1a, and Extended Data Fig. 1). This was most significant for Beta, Delta and Omicron RBDs but was also present, albeit to a lesser extent, for the Gamma variant. Amongst the ACE2bs mAbs, only TAU-2303 maintained its original activity against the Delta strain. TAU-2212 does not bind soluble RBD and could not be evaluated using this assay^9^. To investigate the individual contribution of each mutation, we generated RBDs harboring single or double amino acid substitutions corresponding to variants Beta, Gamma and Delta. Of the eight single-mutated RBDs tested, both the L452R (present in the Delta VOC^23^) and E484K (present in both the Beta and Gamma VOCs^24^) substitutions had a major impact on antibody binding (Figs. 1b and d, and Extended Data Fig. 1). Other single substitutions, however, had no effect on mAb binding. Furthermore, RBDs containing single mutations N439K^25^, Y453F^26,27^ and A475V^28^, which have been reported in some circulating SARS-CoV-2 strains, were bound by all mAbs as strongly as the original wild type RBD. Interestingly, the binding of mAb TAU-2303 to the double mutant K417N / N501Y was reduced although each of the two single mutants separately had no effect (Fig. 1c-d, Extended Data Fig. 1). Overall, we conclude that ACE2bs mAbs are more sensitive to mutations in the RBDs than non-ACE2bs mAbs.

**Fig. 1:**
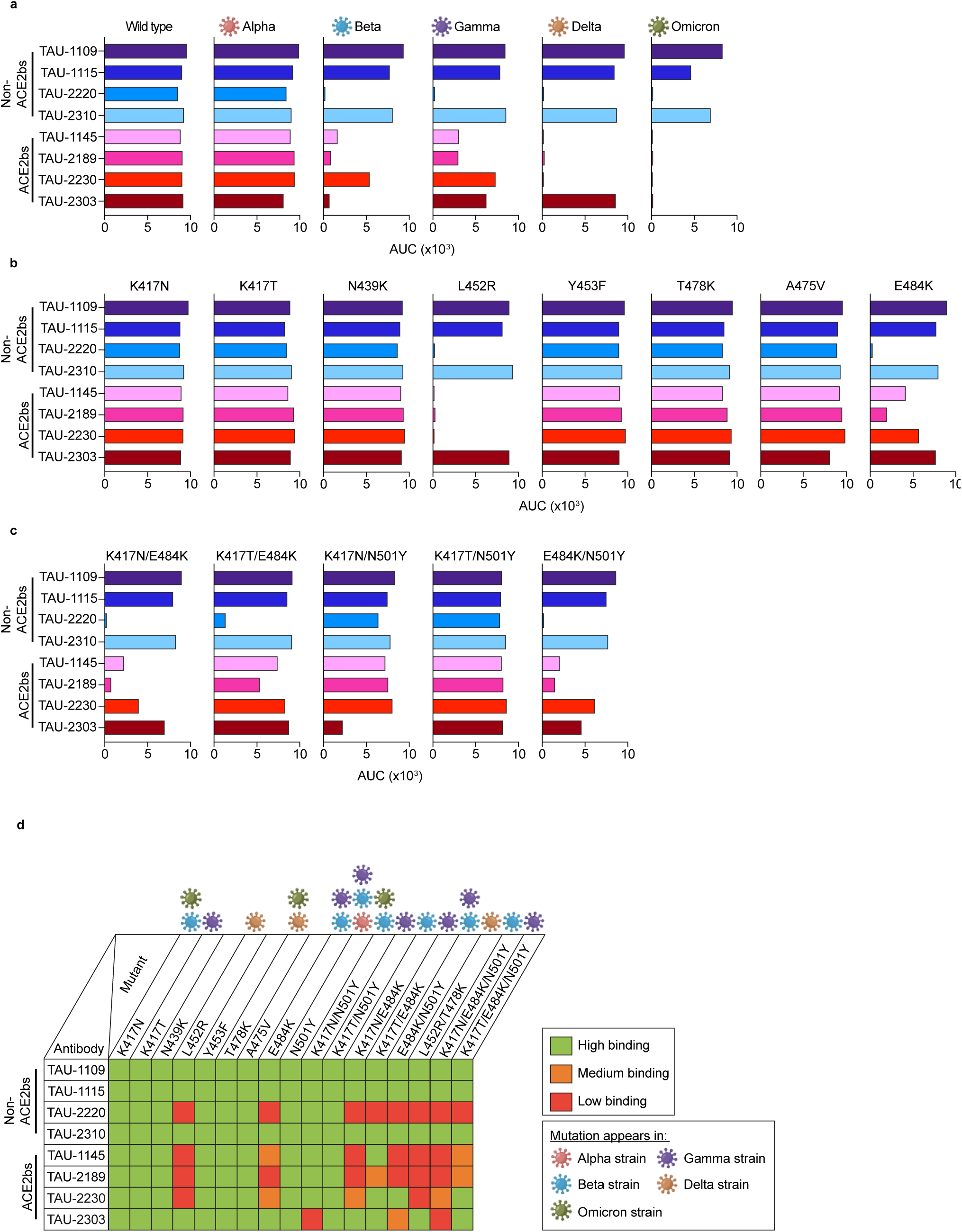
Antibody binding to the RBD of wild type and SARS-CoV-2 VOCs by ELISA. ACE2bs or Non-ACE2bs antibodies are indicated to the left of each panel, **a-d**. For panels **a-c**, each graph represents antibody binding to wild type or VOC (**a**), single (**b**) or double (**c**) mutant RBD. AUC was calculated by GraphPad Prism. The raw OD_650_ values, as well as isotype control, are presented in Extended Data Figure 1. **d**, Summary of the antibody binding affinity to each RBD generated in this study. Green color indicates binding affinity of >75%, orange of 25-75%, and red of <25% as compared to wild type RBD. The VOCs harboring each mutation are indicated on top.

Mutations outside the RBD may also affect the activity of RBD-binding mAbs by altering the conformational organization of the trimer^29^. Therefore, we next expressed the full-length spike protein of wild type, Alpha, Beta, and Delta variants (Extended Data Table S1) on Expi293F cells, and assayed the ability of each mAb to inhibit binding of soluble human ACE2 (hACE2) by flow cytometry. The Gamma full-length spike protein was not produced since the RBD of this variant exhibited similar activity to the Beta RBD. As expected, most of the ACE2bs mAbs effectively inhibited the ACE2:spike_wild type_ and ACE2:spike_Alpha_ interactions, but not those between the ACE2:spike_Beta_ and ACE2:spike_Delta_. The mAb TAU-2212, which can be tested in this assay, demonstrated 25-40% ACE2:spike inhibition when tested against the wild type, Alpha and Delta strains, but had no activity against Beta (Fig. 2a and b, and Extended Data Fig. 2). In fact, none of the ACE2bs mAbs retained their original potency against the Beta variant, and, in accordance with the ELISA data, only mAb TAU-2303 retained its full activity against the Delta variant. As expected, there was no effect in this assay on mAbs TAU-1109, -2310, -1115 and -2220 where the neutralization does not act through receptor blocking.

**Fig. 2:**
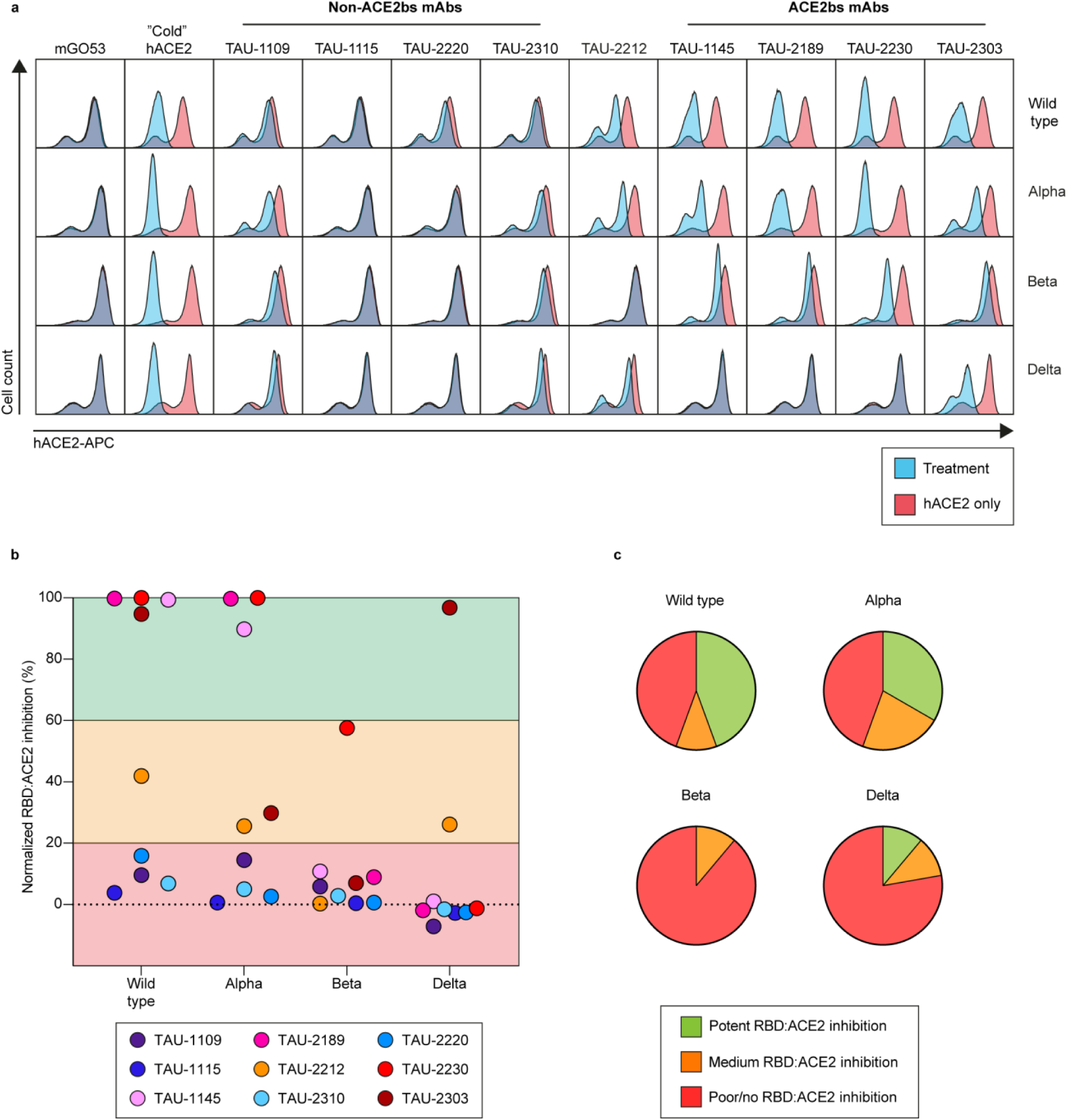
Antibody inhibition of soluble human ACE2 binding to cell expressed spike. **a**, Flow cytometry plots demonstrating the effectiveness of each mAb in interfering with spike:ACE2 binding (see Extended Data Fig. 2 for more details). Expi293F cells were transfected to express the wild type, Alpha, Beta, or Delta spike, and were incubated with each antibody, before being stained with human ACE2 (hACE2) conjugated to APC. Unlabeled hACE2 (“Cold” hACE2) was used as positive control, and mGO53^65^ antibody as an isotype control. The mAbs are indicated on top, and the wild type/VOCs spikes are to the right. Within each plot, the blue histogram indicates treated cells, while the red indicates untreated. **b**, Normalized percent of spike:ACE2 inhibition, calculated by measuring the percent of hACE2-APC positive cells in the presence of each mAb, dividing it by the percent of hACE2-APC positive cells (hACE2 only), and normalizing to 100%. **c**, Pie charts indicating the frequency of spike:ACE2 inhibiting mAbs for wild type and VOCs.

### SARS-CoV-2 VOCs are resistant to most ACE2bs mAbs

We next evaluated the ability of the mAbs to prevent infection. We employed a pseudoviral neutralization assay and an authentic SARS-CoV-2 infection in Vero-TMPRSS2 cells assay to test the activity of the nine mAbs against SARS-CoV-2 wild type and VOCs (Fig. 3). In accordance with the ELISA and flow cytometry results, the Alpha VOC behaved similarly to the wild type strain, while the Beta and Omicron variants were the most resistant, followed by Gamma and Delta. In agreement with the ELISA and flow cytometry results, mAb TAU-2303 was the only ACE2bs mAb that was able to neutralize Delta VOC, with improved potency compared to the wild type strain (Fig. 3a). The non-ACE2bs mAbs TAU-1109 and -2310 retained their efficacy against all tested VOCs, while TAU-2310 exhibited an improved activity against Delta variant compared to the wild type (Fig. 3a and b). These results indicate a crucial support role for non-ACE2bs mAbs in the presence of viral mutations that prevent neutralization by ACE2bs mAbs. Additionally, the improved activity of TAU-2310 and TAU-2303 mAbs demonstrates how genetic variation in a SARS-CoV-2 VOCs can increase neutralization for some classes of mAbs.

**Fig. 3:**
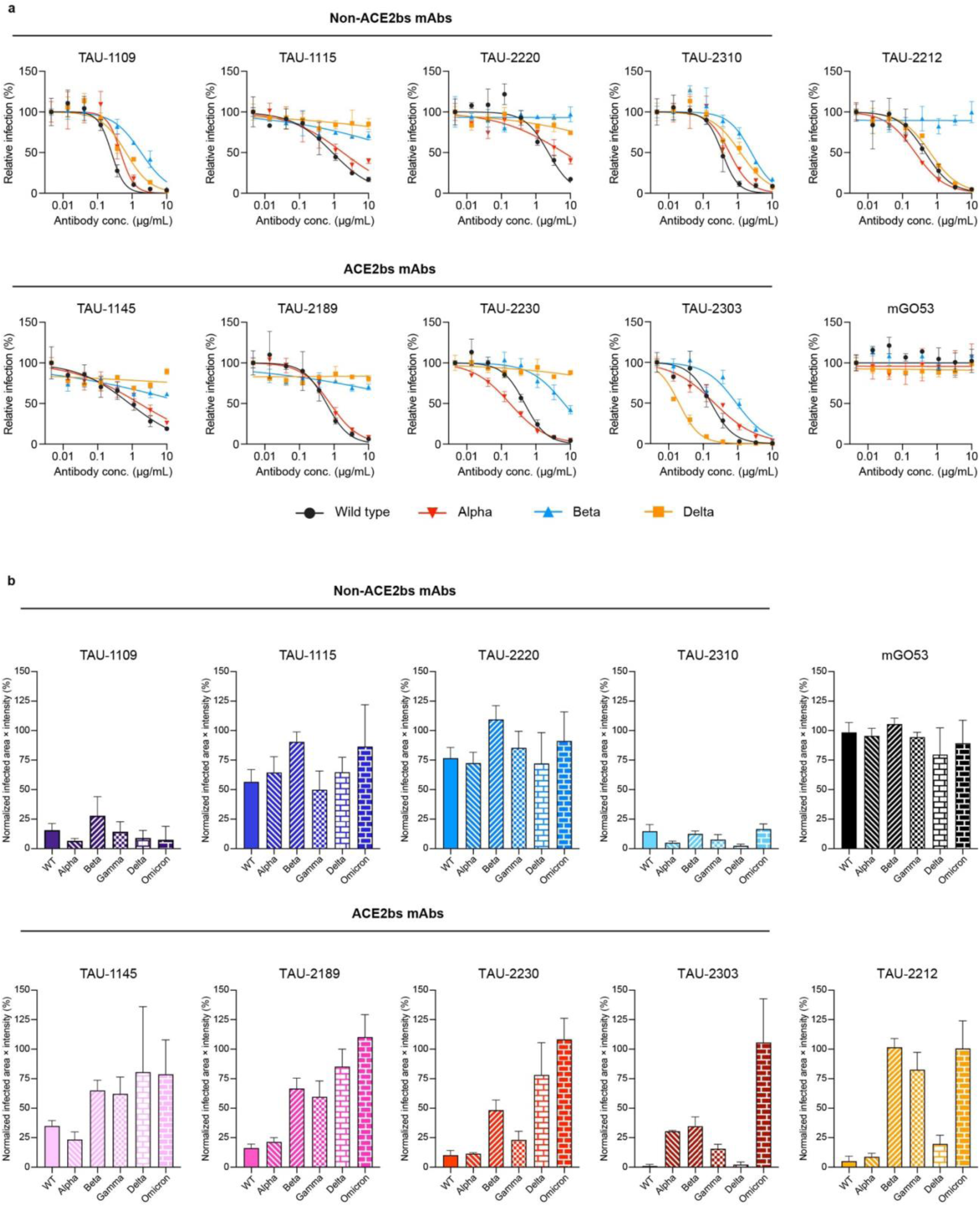
Antibody neutralization of wild type and VOCs. **a**, Neutralization curves of wild type, Alpha, Beta, and Delta VOC pseudo-typed GFP reporter viral particles, by antibodies. For every mAb, each curve represents inhibition of one VOC as indicated below. The mAbs were pre-incubated with the viral particles at 8 consecutive 3-fold dilutions starting at 10 µg/mL of antibody. Fluorescence was read 72 h post infection. Inhibition percentage was calculated by normalizing to untreated cells. Each experiment was done in triplicates. mGO53 was used as an isotype control. **b**, Infected area of Vero-TMPRSS2 by wild type, Alpha, Beta, Gamma, Delta and Omicron authentic SARS-CoV-2. Values were normalized to infected untreated cells. Viral particles were pre-incubated with 100 µg/mL of antibody for 1 h before being added to the cells. Virus infected cells were identified at 24 h post infection using anti-SARS-CoV-2 nucleocapsid AF467-conjugated antibody after cell fixation and permeabilization. Infected cells were quantified using Incucyte S3. mGO53 serves as an isotype control. Statistical analysis was performed using one-way ANOVA and Tukey’s multiple comparison test.

### Structural basis of neutralization of the ACE2bs mAb TAU-2303

Viral evolution appears to be primarily focused on the ACE2bs supersite, consistent with the overall superoir potency of receptor blocking mAbs to inhibit SARS-CoV-2. We decided to focus on mAb TAU-2303, which is the only ACE2bs mAb that is active against the Delta VOC, and mAb TAU-2212 that blocks receptor binding through recognition of a conformational surface, and further investigate the structural basis of these two mAbs. To examine the molecular basis of neutralization mediated by TAU-2303, we first determined the cryoEM structure of the fragment of antigen-binding (Fab) of TAU-2303 (Fab2303) in complex with the ecto SARS-CoV-2 spike trimer, at a resolution of 4.5 Å (Fig. 4a). The cryoEM structure revealed that one Fab2303 molecule binds one spike trimer on the protruding “up” facing RBD. Therefore, mAb TAU-2303 can be categorized as a CoV21^30^-type mAb, that belongs to Class 1 RBD binding mAbs, and binds only one RBD subunit amongst the three available subunits of the trimer (Fig. 4a). We also crystallized Fab2303 in complex with SARS-CoV-2 RBD and analyzed the structure at a resolution of 2.42 Å (Fig. 4b, and Extended Data Table S2). The results from both the crystal and cryoEM structures indicated that Fab2303-RBD complex has a large buried surface of 1185 Å^2^, with the majority of the contact surface (64%) derived from the heavy chain of Fab2303, and only 36% from the light chain. A total of 29 RBD residues in the crystal structure have direct close contacts with Fab2303 (Extended Data Table S3). Consistent with the contact surface analysis, 19 of these residues are from the heavy chain (HC), while 10 are from the light chain (LC) of TAU-2303. The contact residues mediate the formation of a large number of hydrogen bonds (Extended Data Table S3). These include five hydrogen bonds RBD-D420^OD2^ - Fab2303-HC-S56^OG^, RBD-Y473^OH^ - Fab2303-HC-S31^O^, RBD-T415^OG1^ - Fab2303-HC-Y58^OH^, RBD-L455^O^ - Fab2303-HC-Y33^OH^, and RBD-Q493^NE2^ - Fab2303-HC-Y102^OH^ with distances of below 2.7 Å (Extended Data Table S3). In accordance with the biochemistry and in vitro cell assays, 14 (K417, T453, L455, A475, F486, N487, Y489, Q493, G496, Q498, T500, N501, G502, and Y505) of the 29 contacts between Fab2303 and RBD, are also involved in binding the ACE2 receptor, thus confirming that TAU-2303 neutralizes the virus by blocking receptor binding (Figs. 1-3, 4b and Extended Data Figs. 1 and 2). The angle of approach through which Fab2303 binds RBD is 25^°^ relatively to that of ACE2 (Fig. 4b). Further comparisons revealed that the binding mode of mAb TAU-2303 is similar to that of other Class 1 neutralizing antibodies that target RBD (Extended Data Fig. 3)^30-32^.

**Fig. 4.**
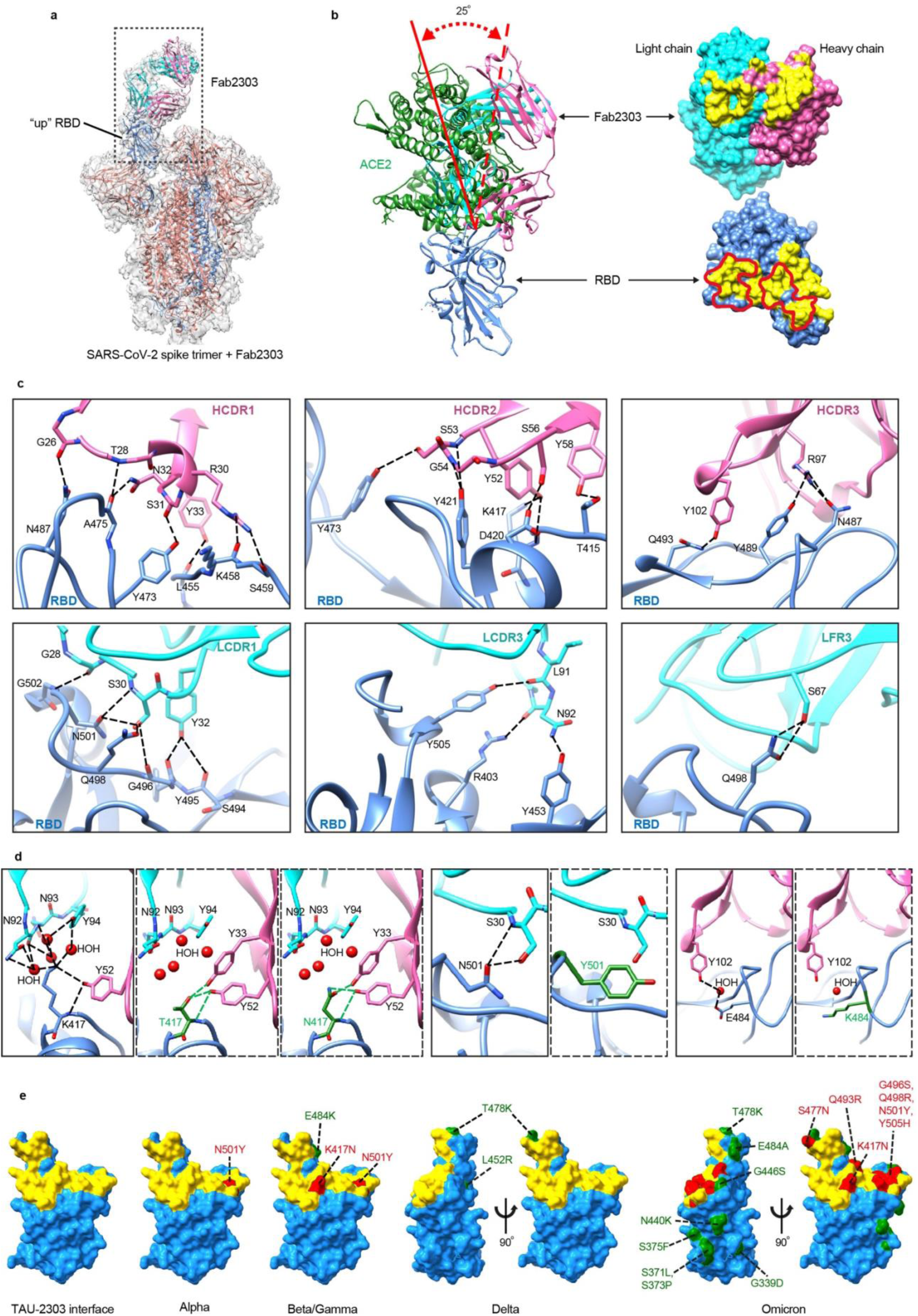
Structural analysis of Fab2303 in complex with the SARS-CoV-2 spike trimer and RBD. **a**, Ribbon diagrams showing the cryoEM structure of the SARS-CoV-2 spike trimer with one Fab2303, binding one of the SARS-CoV-2 spike trimer RBDs in the “up” conformation. The “up” RBD protomer of the spike is colored blue. The other two protomers are colored salmon. The heavy chain of Fab2303 is colored pink and the light chain is colored cyan. **b**, Left: ribbon diagrams showing the Fab2303-RBD crystal structure superimposed onto the ACE2-RBD crystal structure (PDB: 6M0J). The RBD and Fab2303 are colored as in **a**. ACE2 is colored green. The solid and dashed lines in red indicate the long axes of ACE2 and Fab2303, respectively. Right: the paratope and epitope of Fab2303 shown as rendered surface representations. The paratope on Fab2303 and epitope on RBD are colored yellow. Red lines indicate the footprint of ACE2. **c**, Detailed interactions between Fab2303 and SARS-CoV-2 RBD. **d**, Structural comparisons of RBD with RBD mutants K417N, K417T, N501Y, and E484K. Structures of the mutants were modeled in COOT^52^ by using the single mutate function, in which only the side chains of the mutated residues were changed. Most possible side chain conformations of the mutated residues were generated and selected from the rotamer library of COOT and according to the binding energy calculated with PISA. The heavy chain of Fab2303 is colored pink and the light chain is colored cyan. The RBD and Fab2303 are colored as in **a**, with the mutated RBD residues in green. **e**, Surface mapping of key mutations in different variants and the positions of the mutated sites relative to the binding epitopes recognized by mAb TAU-2303. The binding epitopes of TAU-2303 are colored yellow. The mutation sites within or outside the binding epitope of TAU-2303 are colored red and green, respectively.

The ELISA results indicated a reduction in TAU-2303 binding to RBD with the double mutations K417N / N501Y, but not to the protein with the double mutations K417T / N501Y, or any of the single mutations K417N, K417T, or N501Y (Fig. 1 and Extended Data Fig. 1). This agrees with the crystal structure findings that residues K417 and N501 are both in direct contact with TAU-2303. Position 417 is particularly critical for TAU-2303 recognition, as Fab2303-HC-Y52^OH^ forms hydrogen bonds both with the main chain atom N and the side chain atom NH of RBD-K417 (Figs. 4c-d, and Extended Data Table 3). Modeling revealed that when lysine in position 417 is replaced with threonine, both the main chain and side chain hydrogen bonds could be retained (Fig. 4d). Moreover, it becomes possible to establish a new hydrogen bond between the OH atom of the threonine and the OH atom Fab2303-HC-Y33, thus decreasing the total binding energy calculated with PISA^33^ from -9.7 kJ/mol to -9.9 kJ/mol. In contrast, replacing lysine in position 417 by asparagine, increases the length of the hydrogen bonds between the side chain polar atoms ND2H or OD1 with Fab2303-HC-Y33 and Fab2303-HC-Y52, which decreases the favorability of the interaction, as indicated by the increase of the calculated binding energy from -9.7 kJ/mol to -9.5 kJ/mol (Fig. 4d). The asparagine in position 501 has a relatively large number of contacts with Fab2303. The N501 side chain forms two hydrogen bonds with main chain and side chain atoms of Fab2303-LC-S30 (Fig. 4c). While these hydrogen bonds would be completely disrupted by replacing asparagine with tyrosine, our modeling data suggest that new Van der Waals interactions could be formed with tyrosine in this position (Fig. 4d). Since these proposed Van der Waals interactions would not be able to fully compensate for the lost hydrogen bonds, mutant N501Y is predicted to have a slightly lower binding affinity to Fab2303. Interestingly, although glutamate in position 484 was shown to be central to viral escape from neutralizing antibodies^34-36^, our structural analysis indicates that E484 does not make direct interactions with Fab2303. This point was resolved by further examination of the RBD and Fab2303 interface, which revealed that E484 interacts with Fab2303-HC-Y102 through a water molecule (Fig. 4d). The observation that solvent-mediated interactions are usually weaker than direct contacts, is consistent with the results of our functional studies that, in contrast to the other ACE2bs mAbs, the single E484K mutation had no significant impact on TAU-2303 binding (Fig. 1 and Extended Data Fig. 1). The mutations T487K and L452R of RBD_Delta_ are located outside the binding epitope of Fab2303, which, thus explains how TAU-2303 can still neutralize the Delta VOC (Fig. 4e). The structure and modeling data explains why TAU-2303 can bind equally well to an RBD molecule with a single mutation, such as K417N, E484K, and N501Y, but not to an RBD mutant with all three mutations, with the greater cumulative loss of binding energy. Our structural data combining the biochemistry and neutralization results, suggest that the combination of RBD mutations may play a key role in conferring SARS-CoV-2 resistance. Significantly, the Omicron variant, has mutations in seven residues within the binding epitope of Fab2303, including four key contacting residues K417, Q493, Q498, and Y505, providing the structural explanation to the lack of neutralization of Omicron by TAU-2303 (Figs. 3 and 4e).

### mAb TAU-2212 blocks ACE2 binding through conformational dynamics

TAU-2212 mAb is one of the most potent mAbs in our panel, and while not being able to bind soluble RBD by ELISA, it inhibits ACE2 binding as measured by flow cytometry, suggesting that receptor binding is blocked through a different mechanism to that employed by TAU-2303. To investigate the neutralization mechanism of TAU-2212, we prepared Fab2212 by cleaving mAb TAU-2212 with papain. The purified Fab2212 was crystallized and the crystal was diffracted to 2.7 Å. Data analysis revealed that the crystal belongs to the space group P6_5_ with cell dimensions a = 75.98 Å, b = 75.98 Å, and c = 348.14 Å (Extended Data Table 2), and with two molecules in the asymmetric unit. The structure was determined by molecular replacement and was refined to a final R/Rfree of 0.246/0.272. The final model of Fab2212 contains 418 residues, while residues 136-148 and 192-201 of heavy chain are not visible in the map (Fig. 5a). No significant differences were observed between the two molecules in the asymmetric unit (a calculated R.M.S.D. of 0.6 Å between the aligned C_alpha_ atoms, Extended Data Fig. 4). Fab2212 is not able to form a stable complex with the prefusion stabilized SARS-CoV-2 spike ectodomain (Extended Data Fig. 5a), but interacts with spike trimer on the cell surface^9^. With the supposition that mAb TAU-2212 binds a specific conformation of the prefusion spike trimer that is not stable when coated on ELISA plates, we used mAb TAU-2212-protein A coated beads to pull down spike trimers, which were then subjected to cryoEM structural analysis. CryoEM 3-dimensional (3D) classification of the particles revealed that TAU-2212 binds the spike trimer in five distinct conformations (1-5), with 20339, 15868, 14193, 72056, and 39788 particles, respectively (Extended Data Fig. 6 and Extended Data Table 4). In all five conformations, only the Fab portion of mAb TAU-2212 is visible, while the highly flexible Fc region is not seen in the reconstructed maps. Conformations 1, 3 and 4 are composed of two, three and three bound Fabs, respectively, with all the RBDs in the “down” position. In contrast, conformation 2 has one Fab per trimer, with one RBD in the “up” configuration, while the other RBDs that bind the Fab appear in the “down” configuration. Conformation 5 has two head-to-head spike proteins, which are crosslinked by three mAbs (Fig. 5b-c).

**Fig. 5.**
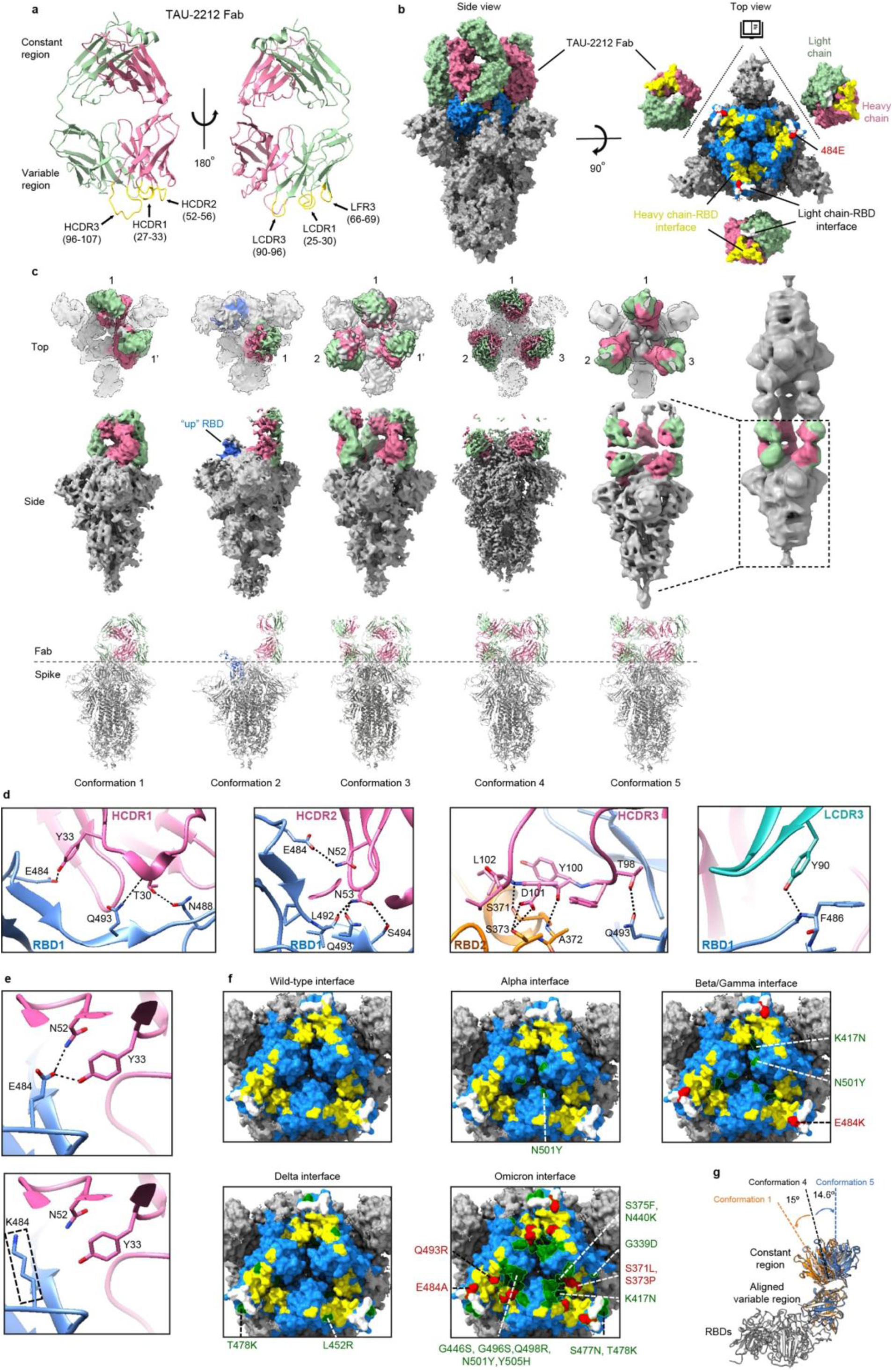
Structural analysis of TAU-2212 in complex with the SARS-CoV-2 spike trimer. **a**, Ribbon diagram showing the crystal structure of Fab2212. The heavy and light chains are colored pink and light green, respectively. The CDR loops are colored yellow. **b**, Surface diagrams showing the side (left) and the top open-up (right) views of the spike trimer with three bound Fabs of mAb TAU-2212. The RBDs of the spike trimer are colored blue and the Fab2212 heavy and light chains are colored as in **a**. The binding epitopes of the heavy and light chains around the junction of two RBDs are colored yellow and white, respectively. Residue E484 is indicated in red. **c**, Surface (top) and ribbon (bottom) diagrams showing the five conformations of the mAb TAU-2212-spike complex. The heavy and light chains of the bound mAbs are colored are colored as in **a**. The spike trimers are in gray. The number pairs X, X’ indicating Fabs from the same mAb. **d**, Detailed interactions between mAb TAU-2212 and the spike trimer illustrated with the high-resolution conformation 1 structure. Heavy chain and light chain CDR loops involved in direct interactions are shown in pink and green, respectively. Residues involved in hydrogen bond formation are shown in sticks with oxygen and nitrogen atoms colored red and blue, respectively. **e**, Ribbon and stick diagrams showing the disruption of the key hydrogen bonds in E484 by the E484K mutation. **f**, Surface diagrams showing mapping of key mutations in different variants and the positions of the mutated sites relative to the binding epitopes recognized by mAb TAU-2212. The TAU-2212 heavy and light chain epitopes are colored yellow and white, respectively. The mutation sites within or outside the binding epitope of TAU-2212 are colored red and green, respectively. **g**, Structural comparisons of the Fabs in conformations 1, 4, and 5. The alignments were performed by using the RBDs (in gray) and the variable regions of the bound Fabs. The bound Fabs of conformations 1, 4, and 5 are colored black, orange, and blue, respectively. Significant differences were observed among the constant regions of the bound Fabs.

We determined the structures of all five conformations to resolutions of 5.54 Å, 7.76 Å, 6.47 Å, 3.45 Å and 9.36 Å (7.32 Å for the split single spike with three Fabs), respectively (Fig. 5b-c, Extended Data Table 4, Extended Data Figs. 6-7). For conformation 4, where the resolution permits ab initio model building and refinement (Extended Data Fig. 8), an atom model of the complexes was built with the crystal structure of Fab2212 and the cryoEM structure of the spike trimer (PDB: 6XEY) as references (Extended Data Table 4). Since the resolution of the maps of conformations 1-3, 5, is too low for ab initio model building, atom models of the complexes were built by fitting the crystal structure of Fab2212 and the cryoEM structure of the S trimer into cryoEM maps.

Conformation 4 of the mAb TAU-2212:spike complex has all the RBDs in the “down” position with three TAU-2212 moieties binding near the junctions between the RBDs. Each Fab2212 binds and crosslinks two adjacent “down” RBDs (RBD1 and 2) with a buried surface of 848 Å^2^ on RBD1 and 292 Å^2^ on RBD2 (Fig. 5b). Comparing the structures of the TAU-2212-bound and free spikes revealed that TAU-2212 binding induces conformational changes in the RBDs, producing a small rotation towards the symmetry axis, and resulting in a more compact RBD head structure after TAU-2212 binding (Extended Data Fig. 9a). In addition, the disordered RBD loops 454-462 and 468-489 become ordered upon binding of TAU-2212 Fabs and have multiple interactions with TAU-2212 (Extended Data Fig. 9b, Extended Data Table 5). These data suggest that TAU-2212 acts by binding and stabilizing the SARS-CoV-2 spike trimer in a “down” orientation and prevents conformational change to “up” RBD which is required for spike:ACE2 interactions.

Most of the epitope recognized by mAb TAU-2212 is on one of the two RBDs (RBD1), and the interactions are predominantly through the heavy chain, especially the loop of complementarity determining region 3 of the heavy chain (HCDR3), which is embedded within the interface of two RBDs. Two CDR loops of the light chain interact directly with four residues on RBD1 (483-486), including the hydrogen bond formed between N atom of F486_RBD1_ and the OH group of Fab2212-HC-Y90 (Fig. 5d). Overall, the interactions between the HCDR loops and RBD1 and RBD2 involve 40 residues, and 14 pairs of hydrogen bonds (Fig. 5d, Extended Data Table 5). These interactions include two hydrogen bonds formed by the OE2 group of E484_RBD1_, with the OH group of Fab2212-HC-Y33, and the ND2 group of Fab2212-HC-N52. The length of the hydrogen bond between the OE2 group of E484_RBD1_ and the OH group of Fab2212-HC-Y33 is 2.5 Å, suggesting that this hydrogen bond could be the main interaction at the contact interface. The mutation E484K completely disrupts the hydrogen bonds between the antibody and the RBD (Fig. 5e and Extended Data Fig. 5b), providing a structural explanation for the complete resistance of the Beta and Gamma VOCs to TAU-2212. Similarly, both the E484A and Q493R mutations present in the new Omicron variant, are considered likely to disrupt the key hydrogen bonds and are therefore expected to affect TAU-2212 binding and reduce neutralization capacity (Fig. 5f). Substitutions S371L and S373P that were reported in the new Omicron variant, also lie within the TAU-2212 interface. To test the effect of these mutations on TAU-2212 binding, we introduced a S373G mutation that did not affect the binding of TAU-2212 (Extended Data Fig. 5b). Taking the effect of all the mutations together, it is therefore not surprising that TAU-2212 does not neutralize Omicron.

Given the flexibility and dynamics of the interaction between TAU-2212 and the spike trimer, we examined binding of the Fab2212 variable regions to the adjacent RBDs in conformations 1-5. We noticed that the densities of bound Fabs2212 were considerably weaker in conformation 2 than in the other conformations, indicating that the “one up” - “two down” configuration may not be favorable for TAU-2212 binding (Fig. 5c). This possibility was also suggested by the pull-down results, where a large portion of the spike protein was in the flowthrough and are likely in the dominant RBD “up” conformations. Structural analysis of the bound Fabs indicated that the constant regions of the three bound Fabs in conformation 4 (three Fabs binding three “down” RBDs), are positioned at a small bending angle (14.6°) relative to the variable regions that are aligned with the z axis (Fig. 5g). In addition, the three molecules are well separated (Fig. 5c) and the distances between the two C254s that form a disulfide bond and crosslink loops of the bound Fabs is approximately 21 Å (Extended Data Fig. 10), suggesting that the three bound Fabs belong to three different mAbs, rather than one mAb binding the trimer with both arms. The constant and variable domains of the bound Fabs in conformation 5 assume similar conformations as these in conformation 4, suggesting that the bound Fabs in conformation 5 belong to three different mAbs. In contrast to conformation 4, only two Fabs are present per trimer in conformation 1. The constant regions of the two bound Fabs in conformation 1 have a large bending angle (29.6°) and are joined at the distal end. In addition, the two C254s that crosslink loops of the bound Fabs have a reasonable distance of 4 Å (Extended Data Fig. 10), suggesting that in this case, the two bound Fabs belong to the same mAb (Fig. 5c and Extended Data Fig. 10). Notably, conformation 3 has the three bound Fabs in two distinct conformations. Two of the Fabs have similar bending angle as that observed in conformation 1, whereas one of the Fabs has similar bending angle as that observed in conformation 5, indicating that the binding of TAU-2212 in conformation 3 is in a mixed mode with two mAbs. Among these, one mAb contributes two Fabs, while one mAb provide only one Fab (Fig. 5c).

Structural analysis of the head-to-head spikes in conformation 5 described three mAbs that crosslink two spikes, with the two Fabs of each mAb in two linearly aligned 180º opposite positions (Fig. 5c and Extended Data Fig. 10c). As a result, the constant region of the bound Fab in conformation 5 has the smallest bending angle and is almost in a plane with the variable region. Given that most particles are in conformation 4, and no spike trimers were observed with all “down” RBDs and a single bound TAU-2212, we can deduce that binding of a single TAU-2212 mAb to the RBD all “down” trimer triggers a conformational change in the RBD that promotes additional Fab binding. The nearby Fab from the same mAb may take the advantage to bind at first. However, the distortion and bending of the bound Fabs from the same mAb make an unstable binding to spike. Thus, the binding mode in conformation 1 could soon be replaced by the binding mode in conformation 4, whereas conformation 3 is likely an intermediate state between conformations 1 and 4. Taken together, mAb TAU-2212 adopts a unique binding mode to spike that could promote quick formation of a multivalent center for aggregates of other spikes or virus particles.

## DISCUSSION

Despite the relatively stable genome of SARS-CoV-2^37^, the continuing spread of the global pandemic has been accompanied by the emergence of new variants with improved transmissibility and mutations that contribute to immune evasion^25,38-41^. With enhanced affinity for human cells and alterations made to vulnerable residues within the spike, these new variants can jeopardies both mAb therapies and vaccines. The Alpha, Beta, Gamma, Delta, and Omicron VOCs are of particular interest as they completely replaced the original Wuhan-Hu-1 strain in subsequent “waves” of the pandemic. The first part of our study assessed the binding inhibition and neutralization of nine antibodies, previously isolated in Israel from Wuhan-Hu-1 SARS-CoV-2 infected individuals^9^. Consistent with other reports^42,43^, we show that ACE2bs mAbs are more affected by viral mutations than mAbs that bind to regions outside the ACE2bs. While these are generally less potent against the original infecting virus, non-ACE2bs mAbs appear to have broader activity against emerging variants.

In the second part of our study, we investigated the mechanism of neutralization of two neutralizing receptor-blocking mAbs, TAU-2303 and TAU-2212, at the atomic level. The atomic structure of Fab2303:RBD confirmed that TAU-2303 mAb binds a surface that is also bound by the ACE2 receptor, as we observed by functional assays. In contrast to other ACE2bs mAbs, TAU-2303 remained active against the Delta VOC, likely due to the comparable modes of binding of ACE2 and TAU-2303 with respect to both epitope and angle of approach to the RBD. However, like the majority of mAbs that block receptor binding, TAU-2303 was less effective against the Beta and completely ineffective against against Omicron variant. We next examined the atypical mAb, TAU-2212, which exhibits an unusual recognition flexibility type of binding involving five different possible conformations. TAU-2212 binds and crosslinks “down” RBDs by displaying an exceptional conformational flexibility. The observation that mAb TAU-2212 binds the spike complex in five different conformations suggests a flexibility of neutralization that is also achieved by stabilization of the spike trimer. The existence of a large number of TAU-2212 mAb crosslinked with head-to-head spikes suggests that TAU-2212 can potently crosslink and aggregate viral particles and thereby reduce the number of effective viral particles in the lung. Furthermore, the binding mode in conformations 1, 3, 4 or 5 would block any “up” conformation of the RBDs, thus, blocking receptor binding. In these ways, TAU-2212 exhibits ACE2bs-like properties while utilizing a distinct mechanism of action. Alternative strategies that broaden neutralization capacity can be deduced from the previously reported antibodies S2M11^44^ and C144^45^ which crosslink two “down” RBDs and have a similar mode of binding to that of TAU-2212 (Extended Data Fig. 11). However, unlike TAU-2212, both S2M11 and C144 can bind the spike trimer with the Fab alone. Out of the three antibodies, S2M11 has the largest binding interface, while the binding interfaces of TAU-2212 and C144 are smaller and more similar (Extended Data Fig. 11). Like TAU-2212, S2M11 mAb promotes a compact RBD head, whereas C144 binding promotes the opposite effect by forcing the RBD head into an open conformation. The larger binding interface of S2M11 encompasses E484 and L452 residues, and therefore is expected to lose efficacy against the Beta, Gamma and Delta variants, whereas TAU-2212 effectively neutralizes Delta (Fig. 3 and Extended Data Fig. 11). Like other mabs in its class TAU-2212 also exhibited a complete loss of activity against the Beta, Gamma and Omicron VOCs.

To summarize, our study provides functional and atomic-level structural data on the interactions between naturally elicited antibodies and SARS-CoV-2 variants. Both TAU-2212 and TAU-2303 exhibit potent neutralization efficacy against the wild type strain and are sensitive to the Omicron variant however, they arise through different B cell developmental programs and are sensitive to different variants and mutations. We therefore conclude that while ACE2bs mAbs can present potent neutralization activity, they are much more sensitive to viral evolution than non-ACE2bs mAbs.

## MATERIALS AND METHODS

### Construction of variant SARS-CoV-2 RBDs

We used our previously reported wild type plasmid^9^ as template for PCR mutagenesis designed to generate RBD constructs harboring single amino acid mutations. Pairs of overlapping DNA primers containing one or two base pair substitutions flanked by 20 bases on each side were designed and synthesized by Syntezza-Israel. PCR reactions were performed using KAPA HiFi HotStart ReadyMix (Roche) DNA polymerase. Each PCR reaction contained 10 µL KAPA HiFi HotStart ReadyMix, 0.5 µM of each primer, and 1 ng template DNA, with the sample volume adjusted to 20 µL with DNase/RNase free water (Bio-Lab). The PCR conditions were as follows: 95 ° C for 3 min, 16 cycles of 98 ° C for 20 s, and 72 ° C for 90 s. Double and triple amino acid mutants were generated similarly with appropriate templates and primers.

### Expression and purification of soluble SARS-CoV-2 RBDs

Each construct was used to transiently transfect Expi293F cells (Thermo Fisher) using the ExpiFectamine 293 Transfection Kit (Thermo Fisher). Seven days post transfection, the cell supernatant was collected, filtered (0.22 µm), and incubated with Ni^2+^-NTA agarose beads (GE Life Sciences) for 2 h at room temperature (RT). Proteins were eluted by 200 mM imidazole, buffer-exchanged to PBS x1, aliquoted, and stored at -80 ° C.

### ELISA

High-binding 96 well ELISA plates (Corning #9018) were coated with 1 µg/mL RBD in PBS x1 overnight at 4 ° C. The following day, the coating was discarded, the wells were washed with “washing buffer” (PBS x1 and 0.05% Tween20) and were blocked for 2 h at RT with 200 µL of “blocking buffer” (PBS x1, 3% BSA (MP Biomedicals), 20 mM EDTA, and 0.05% Tween20 (Sigma)). Antibodies were added at a starting concentration of 4 µg/mL, and seven additional 4-fold dilutions in blocking buffer, and incubated for 1 h at RT. The plates were then washed 3 times with washing buffer before adding secondary anti-IgG (Jackson ImmmunoResearch) antibody conjugated to horseradish peroxidase (HRP) diluted 1:5000 in blocking buffer, and incubation for 1 h at RT. Following four additional washes, 100 µL of TMB (Abcam) was added to each well and the absorbance at 650 nm was read after 20 min (BioTek 800 TS).

### Antibody inhibition of hACE2 binding to cell-expressed spike

Expi293F cells were transfected with pcDNA 3.1 containing SΔC19 of wild type, Alpha, Beta, or Delta variants, using the ExpiFectamine 293 Transfection Kit (Thermo Fisher). The following day, the cells were harvested, centrifuged at 300 x *g*, and resuspended in FACS buffer (PBS x1, 2% FBS and 2mM EDTA). Next, the cells were aliquoted into a 24-well plate (Corning), so that each well contained 3 × 10^6^ cell in 1 mL of FACS buffer. TAU antibodies or mGO53 were added to the appropriate wells at a concentration of 20 µg/mL with unlabeled hACE2 at a concentration of 1 µg/mL. The cells were then incubated for 30 min in an 8% CO_2_ incubator with gentle shaking, transferred to FACS tubes, washed with FACS buffer, and incubated with biotinylated hACE2 for 20 min at 4 ° C. Following an additional washing step, the cells were incubated with streptavidin-APC (Miltenyi Biotec, 130-106-792) and washed again. APC florescence was recorded using a CytoFLEX S4 (Beckman Coulter).

### Pseudo-particle preparation and neutralization assays

SARS-CoV-2-spike pseudo-particles were obtained by co-transfecting Expi293F cells with pCMV delta R8.2, pLenti-GFP (Genecopoeia), and pcDNA 3.1 SΔC19 (Thermo Fisher) at a ratio of 1:2:1, respectively, according to the manufacturer’s instructions. The supernatant was harvested 72 h post transfection, centrifuged at 1500 x *g* for 10 min and passed through a 0.45 μm filter (LIFEGENE, Israel). The supernatant was then concentrated to 5% of its original volume using an Amicon Ultra with a 100 KDa cutoff at 16 °C (Merck Millipore). HEK-293 cells stably expressing hACE2 were seeded into 0.1% gelatin-coated 96-well plates (Greiner) at an initial density of 0.75 × 10^5^ cells per well. The following day, concentrated pseudo-particles were incubated with serial dilutions of antibodies for 1 h at 37 °C and added to the 96 well plates. After 48 h, the cell medium was replaced with fresh DMEM medium excluding Phenol Red, and 24 h later, the 96-well plates were imaged by IncuCyte ZOOM (Essen BioScience). Cells were imaged with a 10x objective using the default IncuCyte software settings, which were used to calculate number of GFP-positive cells from four 488 nm-channel images in each well (data for each antibody was collected in triplicates). The number of GFP-positive cells was normalized and converted to a neutralization percentage. pseudo-particles expressing the Alpha, Beta and Delta spikes were produced and tested similarly (Extended Data Table 1).

### Virus preparation and titer determination

All work with SARS-CoV-2 was conducted under Biosafety Level-3 conditions at the University of California San Diego. SARS-CoV-2 isolates WA1 (USA-WA1/2020, NR-52281), Beta (B.1.351, hCoV-19/South Africa/KRISP-K005325/2020, NR-54009), Gamma (P.1, hCoV-19/Japan/TY7-503/2021, NR-54982), and Delta (B.1.617.2, hCoV-19/USA/PHC658/2021, NR-55611) were acquired from BEI Resources. Viral stocks originally isolated on VeroE6 were passaged once through primary human bronchial epithelial cells (NHBECs) differentiated at air-liquid interface (ALI) before expansion on VeroE6-TMPRSS2 (Sekisui XenoTech), referred to here as Vero-TMPRSS2. Variant Alpha (B.1.1.7) was isolated on NHBECs at ALI from a nasopharyngeal swab and expanded on Vero-TMPRSS2. Isolate has been deposited at BEI Resources as hCoV-19/USA/CA_UCSD_5574/2020, NR-54020. All viral stocks were verified by deep sequencing.

Virus titers were validated using a combination of fluorescent focus assay and tissue culture infectious dose (TCID)_50_ assays on Vero-TMPRSS2 and Calu-3 (ATCC) monolayers. Serial dilutions of virus stocks in DMEM (Corning, #10-014-CV) were added to Vero-TMPRSS2 monolayers in 96-well plates and incubated for 1 h at 37 °C with rocking. For fluorescent focus assays, cells were overlaid with 1% methylcellulose in DMEM supplemented wth 2% FBS and 1x Pen/Strep. Plates were incubated at 37 °C in 5% CO_2_ for 24h depending on the assay and then fixed with a final concentration of 4.5% formaldehyde for at least 30 min at RT. For TCID_50_ assays, Vero-TMPRSS2 or Calu3 cells were infected as above and 100µL DMEM or MEM with 2% FBS was subsequently added per well. Monolayers were observed for at least 4 days for appearance of CPE and then fixed as above and stained with antibody against SARS-CoV-2 Nucleocapsid. TCID_50_ was calculated using the Reed-Muench method^46-48^.

### Immunofluorescence imaging and analysis

For viral nucleocapsid detection in Vero-TMPRSS2 cells by immunofluorescence, cells were washed twice with PBS x1 and fixed in 4% formaldehyde for 30 min at RT. Fixed cells were washed with PBS x1 and permeabilized for immunofluorescence using BD Cytofix/Cytoperm according to the manufacturer’s protocol for fixed cells, and then stained for SARS-CoV-2 with a primary nucleocapsid antibody (GeneTex GTX135357) and a secondary anti-rabbit AF647 antibody (ThermoFisher A20185), The nuclei were counterstained with Sytox Green. Infected cells from whole well scans were identified using the Incucyte S3. Data were logged from the Incucyte analysis modules and graphed with GraphPad Prism 8.

### Expression and purification of soluble RBD, spike and antibodies for crystallization

SARS-CoV-2 RBD (residues Arg319 to Lys529) was expressed by using the Bac-to-Bac Baculovirus System (Invitrogen). RBD containing the gp67 signal peptide and a C-terminal 6×His tag was inserted into pFastBac1 to form the plasmid pFastBac1-RBD. The plasmid was then transformed into DH10 Bac component cells. The recombinant bacmid was extracted and further transfected into Sf9 cells using Cellfectin II (Invitrogen, #10362100). The recombinant viruses were harvested from the transfected supernatant and amplified to generate high-titer virus stock. The viruses were then used to infect Hi5 cells for RBD expression. Secreted RBD in the supernatant was harvested and applied to cobalt agarose beads, then eluted with 300 mM imidazole, and further purified by using a Superdex 200 Increase 10/300 column (GE Healthcare) running in a buffer containing 20 mM HEPES at pH 8.0 and 150 mM NaCl.

The heavy chain and light chain of TAU-2303 or TAU-2212 were cloned separately into a pCMV vector and transiently transfected into HEK293F cells by using PEI at a ratio of 1:1. Supernatant was harvested four days after transfection. TAU-2212 in the supernatant was captured and purified by using protein A beads (GE healthcare), and was eluted by 0.1 M glycine at pH 3.2, then neutralized to pH 7.6.

For Fab preparation, the purified antibodies TAU-2303 and TAU-2212 were digested by using the protease papain (Sigma, #P3125) with an IgG to papain ratio of 66:1 (w/w) for 3 hours at 37 °C. The undigested antibodies were removed with Protein A Sepharose (GE Healthcare) and the flow through was collected, and further purified by using a Superdex 200 Increase 10/300 column (GE Healthcare) running in a buffer containing 20 mM HEPES at pH 8.0 and 150 mM NaCl.

The extracellular domain of S protein (S-ECD) (1-1208 amino acids, Genebank ID: QHD43416.1) was cloned into the pCMV vector with six proline substitutions at residues 817, 892, 899, 942, 986, and 987 (S6P mutant) or two proline substitutions at residues 967 and 968 (S2P mutant)^49^, a “GSAS” substitution at residues 682 to 685, and a C-terminal T4 fibritin trimerization motif followed by a StrepII tag (S6P mutant) or a Flag tag (S2P mutant). The S2P or S6P pCMV plasmids were used to transiently transfect HEK293F cells with polyethylenimine (PEI) (Polysciences, #24765). The recombinant S6P was affinity purified from the cell supernatant by using StrepTactin resin (IBA). The eluted material was further purified by size-exclusion chromatography using a Superose 6 10/300 column (GE Healthcare) running in 20 mM HEPES pH 8.0 and 150 mM NaCl. Supernatant containing S2P was collected four days after transfection, S2P was affinity purified by anti-Flag antibody beads and was eluted by 0.1 mg/mL 3×Flag peptide in 20 mM HEPES at pH 7.6 and 150 mM NaCl.

### Complex preparation and crystallization

The Fab portion of TAU-2303 was mixed with RBD at a molar ratio of 1:2 at 4 °C, and the resultant complex was purified by size-exclusion chromatography with a Superdex 200 Increase 10/300 column running in 150 mM NaCl and 20 mM Tris-HCl, pH 8.0. The peak fractions containing the complex were collected and concentrated to 6.5 mg/mL for crystallization. The RBD-Fab2303 crystals were grown at 18 °C by using the hanging-drop vapor diffusion method with 1 μL protein mixed with 1 μL of reservoir solution containing 0.2 M sodium tartrate dibasic dihydrate and 14% (w/v) polyethylene glycol 3350. Crystals were soaked in the reservoir solution supplemented with 15% glycerol, and flash-frozen in liquid nitrogen for data collection.

The Fab portion of TAU-2212 was concentrated to 7 mg/mL for crystallization. The crystal was grown at 16 °C in 26% (w/v) polyethylene glycol 3350, 0.2 M (NH_4_)_2_SO_4_, pH 8.0. Crystals were soaked in the reservoir solution supplemented with 10% glycerol, and flash-frozen in liquid nitrogen for data collection.

### Data collection, structure determination, and refinement

The X-ray diffraction data were collected at the beamlines BL18U (RBD-Fab2303) and BL02U (Fab2212) of the Shanghai Synchrotron Research Facility. Data were processed and scaled with HKL2000^50^. The structure was determined by molecular replacement using PHASER^51^. Manual building and adjustments of the structures were performed in COOT^52^. The structure of RBD-Fab2303 was refined by using PHENIX^53^. Data processing showed that the crystal of Fab2212 belongs to the space group P6_5_. However, the merohedral twinning appears to assign the crystal to the space group P6_5_22. The structure of Fab2212 was refined against the twinned data by using Refmac5^54^ with the twinning operator k, h, -l and an estimated initial twin ratio of 0.52 to 0.48 between the two domains. The final refined twin ratio between the two domains were 0.59 to 0.41. Data collection and refinement statistics are listed in Extended Data Table 2. Structural analysis of antibody–antigen contacts were assessed through CCP4i^55^ (Extended Data Table 2 and 5). All structural representations were prepared through the use of the UCSF Chimera and ChimeraX^56,57^.

### CryoEM sample preparation, data acquisition, and processing

Fab 2303 was mixed with the purified S6P trimer (2:1 molar ratio Fab per protomer) to form the Fab–S complex at a concentration of 2 mg/mL. The mixture was incubated on ice for 30 minutes. then, 3 μL of the mixture was applied to a glow-discharge holey carbon grids (Quantifoil, Cu 200 mesh, R1.2/1.3). The grid was blotted for 4.5 s in 100% humidity before being flash frozen in liquid ethane by using a Vitrobot Mark IV (Thermo Fisher). For TAU-2212-S2P complex preparation, S2P and TAU-2212 were incubated for 30 min on ice at a molar ratio of 1:2. The antibody-spike complex was purified by protein A beads. The eluted complex was crosslinked by 0.125% glutaraldehyde for 20 min. The crosslinked sample was purified by size-exclusion chromatography with a Superose 6 10/300 column. The target fraction was collected and concentrated to 0.8 mg/ml for cryoEM grid preparation by using similar conditions to those used for S6P and Fab2303.

Image data of the S6P-Fab2303 and S2P-mAb-TAU2212 complexes were collected on a Titan Krios electron microscope (FEI Company) equipped with a field emission gun operated at 300 kV and a Gatan K3 Summit camera. Images of S6P-Fab2303 were recorded at a defocus range of -1.5 to -2.8 μm, with a pixel size of 1.25 Å. A total dose of ∼ 50 electrons per Å^2^ was used. SerialEM was used for the data collection. A total of 2599 movie stacks for S6P-Fab 2303 and 4675 movie stacks for S2P-TAU2212 were collected. The frames in each movie stack were aligned, summed, and 2× binned by using the program Motion Cor2^58^. The CTF parameters of the micrographs were determined by using the program Gctf with local defocus variations taken into consideration^59^. For the S6P-Fab2303 complex, a total of 546293 particles were boxed by using Gautomatch, and were then subjected to 2D classification using RELION^60^. The cryoEM structure of the closed state SARS-CoV-2 spike (PDB accession number: 6VXX^61^) was low-pass filtered to 40 Å and used as the initial model. A total of 38331 particles were selected for the final 3D refinement without imposing any symmetry, which yielded a cryoEM map at a resolution of 4.5 Å (Extended Data Table 4).

For the S2P-mAb-TAU2212 complex, data was collected at defocus -1.5 to -2.0 µm, with pixel size of 0.97Å. 90569 particles were picked by Gautomatch and then were subjected to 2D classifications with RELION 3.0. The 2D classification results showed two major classes, one with a single spike and the other with two head-to-head crosslinked spikes. Particles in the two classes were split for separated 3D classifications in RELION 3.0. For the head-to-head crosslinked spikes, 39788 particles were selected for 3D refinement with D3 symmetry imposed, which resulted in a cryoEM map at a resolution of 9.36 Å. To improve the reconstruction, the head-to-head spike particles were split into two particles each with one spike and three bound Fabs. The split particles were subjected to local refinements, which resulted in a cryoEM map at a resolution of 7.3 Å (Extended Data Fig. 6).

Particles in classes with a single spike, were selected and subjected to 3D classification with a S trimer cryoEM map as the reference (PDB accession number: 6XEY^62^). This produced four distinct conformations: conformation 1 has two bound Fabs; while conformation 2 has only one bound Fab; conformation3 has two fabs from one mAb and one fab from other mAb; conformation 4 is 3-fold symmetric and has three bound Fabs. Particles from each conformation were selected and used for refinement against an RBD “all-down” spike that was low passed to 60 Å. For conformation 4, the final refinements were performed with 72056 selected particles and C3 symmetry imposed. The resolution of the final reconstructed map is 3.45 Å. Side chain densities are clearly visible in most of the reconstructed map (Extended Data Fig. 8). For conformations 1, 2 and 3, the final refinements were performed with 20339, 15868 and 14193 selected particles, respectively, and without imposing any symmetry. The refinements resulted in a cryoEM map at a resolution of 5.54 Å for conformation 1, map at a resolution of 7.76 Å for conformation 2, and a cryoEM map at a resolution of 6.47 Å for conformation 3.

All of the refined density maps were applied with a negative B-factor and corrected for the modulation transfer function (MTF) of the detector. The reported resolution is based on the gold-standard Fourier shell correlation (FSC) 0.143 criterion^63,64^ (Extended Data Fig. 7).

## Acknowledgments

We thank the members of the Xiang and Freund labs for fruitful discussions and assistance. NTF is funded by the ISF grant number #1422/18. MGT and NTF are funded by KillCorona ISF grant number #3711/20. YX is funded by the Spring Breeze Fund of Tsinghua University, the National Natural Science Foundation of China (grants: 31925023, 21827810, 31861143027), the Ministry of Science and Technology of China (grant 2021YFA1300200), the Beijing Frontier Research Center for Biological Structure, and the Beijing Advanced Innovation Center for Structure Biology. BA is supported by NIH grant (K08 AI130381) and AFC is supported by Career Award for Medical Scientists from the Burroughs Wellcome Fund.

## Author contributions

MM planned and performed the biochemical experiments, analysed data, prepared the figures, and wrote the manuscript together with NTF and YX and BAC. RFL and BTM performed crystal and cryoEM structure determination and analysis, prepared the figures, and wrote the manuscript together with NTF and YX and BAC. MW, JA MD and MGT preformed the pseudo-viral assays. JCL performed cell line maintenance and plate setup for Vero assays AEC and AFC conducted virus generation and tittering, SLL helped with BSL-3 with assay optimization. YX, NTF and BAC planned and supervised the experiments, analyzed the data and wrote the manuscript.

## Data and materials availability

The atomic coordinates and EM maps have been deposited into the Protein Data Bank (http://www.pdb.org) and the EM Data Bank, respectively: Fab2303-RBD complex (PDB-####, ####), Fab2303-S complex (EMD-####, ####), Fab2212 (PDB-####), mAb2212-S complex (EMD-####, ####, ####, ####, ####, ####).

## Supplementary figures legends

**Extended Data Fig. 1: Antibody binding to wild type SARS-CoV-2 and VOCs RBDs by ELISA**. Representative ELISA curves for the eight RBD-binding mAbs against (indicated on the bottom) to wild type RBD and RBDs harboring single, double or triple mutations corresponding to common SARS-CoV-2 VOCs. The y axis shows OD_650_ values, and the x axis shows Log [mAb concentration], starting with 10 µg/mL, and with 8 consecutive 4-fold dilutions. mGO53 was used as an isotype control antibody.

**Extended Data Fig. 2: Inhibition of hACE2 binding to cell expressed spike by antibodies as measured by flow cytometry. a**, Gating strategy used to measure antibody inhibition of spike:ACE2 binding. **b-e**, Representative flow cytometry plots of positive hACE2 cells expressing wild type (**b**), Alpha (**c**), Beta (**d**) and Delta (**e**) spikes in the presence of mAbs. mGO53 was used as an isotype control. Unlabeled (“cold” hACE2) was used as positive control for inhibition.

**Extended Data Fig. 3: Comparison between the epitope of TAU-2303 and other Class 1 mAbs. a**, Ribbon diagrams showing the comparisons of binding directions to RBD among Fab2303 (yellow), BG4-25^31^ (green), C098^32^ (pink) and C099^32^ (orange). The Fab-RBD complexes are superimposed with the RBD (blue). **b**, Comparisons of contact interfaces on RBD. The surface of RBDs are colored blue and the binding epitopes are in yellow.

**Extended Data Fig. 4: Crystal structure of Fab2212. a**, Ribbon diagrams showing two Fab2212 molecules in the asymmetric unit of the crystal. **b**, Superimposition of the two Fab2212 molecules in the asymmetric unit of the crystal.

**Extended Data Fig. 5: Binding of mAb TAU-2212 or Fab TAU-2212 to spike trimer. a**, SDS-PAGE gel analysis showing the pull-down of mAb TAU-2212 or Fab TAU-2221 by S2P. **b**, SDS-PAGE gel analysis showing the pull-downs of S2P or S2P mutants by mAb TAU-2212.

**Extended Data Fig. 6: Cryo-EM data processing workflow for the complex of TAU-2212 and spike trimer**.

**Extended Data Fig. 7: Orientation distributions, FSC curves and local resolution maps of the reconstructions of the five TAU-2212-S2P conformations. a**. The orientation distributions of particles used for final refinements of each reconstruction. **b**. Fourier shell coefficient curves of the reconstructions. The FSC threshold is 0.143. **c**. Local resolution maps calculated by ResMap with the density maps^66^.

**Extended Data Fig. 9: Structural comparisons of TAU-2212 bound and free spike trimers. a**, Top panel: ribbon diagrams showing one protomer of the TAU-2212 bound spike trimer superimposed on one protomer of the TAU-2212 free spike trimer. Superimpositions of the structures were performed with the S2 portion only. Bottom panel: ribbon diagrams and the long axes of the aligned RBDs showing the TAU-2212 binding induced conformational changes. **b**, Ribbon diagrams showing that loops 454-462 and 468-489 become ordered upon TAU-2212 binding.

**Extended Data Fig. 10: Structural modeling of the spike trimer bound by full length TAU-2212. a**, Sequence alignment between the TAU-2212 heavy chain and the human IgG B12 heavy chain^67^. The flexible loop “DKTHT” that connects the Fc and Fab of the antibody is highlighted in lime and the cysteines that form a disulfide bond between the loops are highlighted in yellow. **b**. Ribbon diagram of the crystal structure of human IgG B12, PDB ID:1HZH. Heavy chains are in green and light chains are in salmon. The disulfide bonds are in yellow and the flexible loops are colored lime. **c**. B12 Fab fitted in the reconstructed maps of conformations 1, 4, 5. The constant and variable regions of the Fab were fitted separately as rigid bodies. Zoom in ribbon diagrams exhibit the distances between two fitted Fabs. The distance between the C_alpha_ atoms of the two C254s was measured and indicated.

**Extended Data Fig. 11: Epitopes of antibodies similar to TAU-2212**. Contact epitopes of S2M11 (**a**) and C144 (**b**) are shown in yellow. Key mutation sites within or outside the binding epitope of TAU-2303 are colored red and green, respectively.

**Extended Data Table 1: Spike and RBD mutations of the Alpha, Beta, Gamma, Delta and Omicron VOCs**.

**Extended Data Table 2: X-ray data collection and refinement statistics for RBD-Fab2303 complex and Fab2212, related to Figure 4 and 5**.

**Extended Data Table 3: Contacts between RBD and Fab2303**.

**Extended Data Table 4: Cryo-EM data collection, refinement and validation statistics for RBD-Fab2303 and spike-TAU-2212**.

**Extended Data Table 5: Contacts between spike and TAU-2212**.

## References

1 Garcia-Beltran, W. F. et al. Multiple SARS-CoV-2 variants escape neutralization by vaccine-induced humoral immunity. Cell 184, 2523, doi:10.1016/j.cell.2021.04.006 (2021).

2 Lucas, C. et al. Impact of circulating SARS-CoV-2 variants on mRNA vaccine-induced immunity. Nature, doi:10.1038/s41586-021-04085-y (2021).

3 Bates, T. A. et al. Neutralization of SARS-CoV-2 variants by convalescent and BNT162b2 vaccinated serum. Nat Commun 12, 5135, doi:10.1038/s41467-021-25479-6 (2021).

4 Brouwer, P. J. M. et al. Potent neutralizing antibodies from COVID-19 patients define multiple targets of vulnerability. Science 369, 643–650, doi:10.1126/science.abc5902 (2020).

5 Kreer, C. et al. Longitudinal Isolation of Potent Near-Germline SARS-CoV-2-Neutralizing Antibodies from COVID-19 Patients. Cell 182, 843–854 e812, doi:10.1016/j.cell.2020.06.044 (2020).

6 Robbiani, D. F. et al. Convergent antibody responses to SARS-CoV-2 in convalescent individuals. Nature, doi:10.1038/s41586-020-2456-9 (2020).

7 Rogers, T. F. et al. Isolation of potent SARS-CoV-2 neutralizing antibodies and protection from disease in a small animal model. Science, doi:10.1126/science.abc7520 (2020).

8 Wec, A. Z. et al. Broad neutralization of SARS-related viruses by human monoclonal antibodies. Science, doi:10.1126/science.abc7424 (2020).

9 Mor, M. et al. Multi-clonal SARS-CoV-2 neutralization by antibodies isolated from severe COVID-19 convalescent donors. PLoS Pathog 17, e1009165, doi:10.1371/journal.ppat.1009165 (2021).

10 Hansen, J. et al. Studies in humanized mice and convalescent humans yield a SARS-CoV-2 antibody cocktail. Science 369, 1010–1014, doi:10.1126/science.abd0827 (2020).

11 Baum, A. et al. Antibody cocktail to SARS-CoV-2 spike protein prevents rapid mutational escape seen with individual antibodies. Science 369, 1014–1018, doi:10.1126/science.abd0831 (2020).

12 Chen, P. et al. SARS-CoV-2 Neutralizing Antibody LY-CoV555 in Outpatients with Covid-19. N Engl J Med 384, 229–237, doi:10.1056/NEJMoa2029849 (2021).

13 Chen, R. E. et al. In vivo monoclonal antibody efficacy against SARS-CoV-2 variant strains. Nature 596, 103–108, doi:10.1038/s41586-021-03720-y (2021).

14 Wang, P. et al. Antibody resistance of SARS-CoV-2 variants B.1.351 and B.1.1.7. Nature 593, 130–135, doi:10.1038/s41586-021-03398-2 (2021).

15 Hoffmann, M. et al. SARS-CoV-2 variants B.1.351 and P.1 escape from neutralizing antibodies. Cell 184, 2384–2393 e2312, doi:10.1016/j.cell.2021.03.036 (2021).

16 Copin, R. et al. The monoclonal antibody combination REGEN-COV protects against SARS-CoV-2 mutational escape in preclinical and human studies. Cell 184, 3949–3961 e3911, doi:10.1016/j.cell.2021.06.002 (2021).

17 Rambaut, A. et al. Preliminary genomic characterisation of an emergent SARS-CoV-2 lineage in the UK defined by a novel set of spike mutations. https://virological.org/t/preliminary-genomic-characterisation-of-an-emergent-sars-cov-2-lineage-in-the-uk-defined-by-a-novel-set-of-spike-mutations/563 (2020).

18 Tegally, H. et al. Detection of a SARS-CoV-2 variant of concern in South Africa. Nature 592, 438–443, doi:10.1038/s41586-021-03402-9 (2021).

19 Faria, N. R. et al. Genomic characterisation of an emergent SARS-CoV-2 lineage in Manaus: preliminary findings. https://virological.org/t/genomic-characterisation-of-an-emergent-sars-cov-2-lineage-in-manaus-preliminary-findings/586 (2021).

20 Mlcochova, P. et al. SARS-CoV-2 B.1.617.2 Delta variant replication and immune evasion. Nature, doi:10.1038/s41586-021-03944-y (2021).

21 Edara, V. V. et al. Infection and Vaccine-Induced Neutralizing-Antibody Responses to the SARS-CoV-2 B.1.617 Variants. N Engl J Med 385, 664–666, doi:10.1056/NEJMc2107799 (2021).

22 Gowrisankar, A., Priyanka, T. M. C. & Banerjee, S. Omicron: a mysterious variant of concern. Eur Phys J Plus 137, 100, doi:10.1140/epjp/s13360-021-02321-y (2022).

23 Deng, X. et al. Transmission, infectivity, and neutralization of a spike L452R SARS-CoV-2 variant. Cell 184, 3426–3437 e3428, doi:10.1016/j.cell.2021.04.025 (2021).

24 Weisblum, Y. et al. Escape from neutralizing antibodies by SARS-CoV-2 spike protein variants. Elife 9, doi:10.7554/eLife.61312 (2020).

25 Thomson, E. C. et al. Circulating SARS-CoV-2 spike N439K variants maintain fitness while evading antibody-mediated immunity. Cell 184, 1171–1187 e1120, doi:10.1016/j.cell.2021.01.037 (2021).

26 Fonager, J. et al. Working paper on SARS-CoV-2 spike mutations arising in Danish mink, their spread to humans and neutralization data. https://files.ssi.dk/Mink-cluster-5-short-report_AFO2 (2020).

27 ECDC. Detection of new SARS-CoV-2 Variants Related to Mink. https://www.ecdc.europa.eu/sites/default/files/documents/RRA-SARS-CoV-2-in-mink-12-nov-2020.pdf (2020).

28 Li, Q. et al. The Impact of Mutations in SARS-CoV-2 Spike on Viral Infectivity and Antigenicity. Cell 182, 1284–1294 e1289, doi:10.1016/j.cell.2020.07.012 (2020).

29 Mansbach, R. A. et al. The SARS-CoV-2 Spike variant D614G favors an open conformational state. Sci Adv 7, doi:10.1126/sciadv.abf3671 (2021).

30 Barnes, C. O. et al. Structures of Human Antibodies Bound to SARS-CoV-2 Spike Reveal Common Epitopes and Recurrent Features of Antibodies. Cell, doi:10.1016/j.cell.2020.06.025 (2020).

31 Scheid, J. F. et al. B cell genomics behind cross-neutralization of SARS-CoV-2 variants and SARS-CoV. Cell 184, 3205–3221 e3224, doi:10.1016/j.cell.2021.04.032 (2021).

32 Muecksch, F. et al. Affinity maturation of SARS-CoV-2 neutralizing antibodies confers potency, breadth, and resilience to viral escape mutations. Immunity 54, 1853–1868 e1857, doi:10.1016/j.immuni.2021.07.008 (2021).

33 Potterton, L. et al. CCP4i2: the new graphical user interface to the CCP4 program suite. Acta Crystallogr D Struct Biol 74, 68–84, doi:10.1107/S2059798317016035 (2018).

34 Greaney, A. J. et al. Mapping mutations to the SARS-CoV-2 RBD that escape binding by different classes of antibodies. Nature Communications 12, doi:ARTN 4196 10.1038/s41467-021-24435-8 (2021).

35 Alenquer, M. et al. Signatures in SARS-CoV-2 spike protein conferring escape to neutralizing antibodies. Plos Pathogens 17, doi:ARTN e1009772 10.1371/journal.ppat.1009772 (2021).

36 Wang, L. S. et al. Ultrapotent antibodies against diverse and highly transmissible SARS-CoV-2 variants. Science 373, 759-+, doi:10.1126/science.abh1766 (2021).

37 Dearlove, B. et al. A SARS-CoV-2 vaccine candidate would likely match all currently circulating variants. P Natl Acad Sci USA 117, 23652–23662, doi:10.1073/pnas.2008281117 (2020).

38 Tao, K. M. et al. The biological and clinical significance of emerging SARS-CoV-2 variants. Nat Rev Genet 22, 757–773, doi:10.1038/s41576-021-00408-x (2021).

39 Boehm, E. et al. Novel SARS-CoV-2 variants: the pandemics within the pandemic. Clin Microbiol Infec 27, 1109–1117, doi:10.1016/j.cmi.2021.05.022 (2021).

40 Karim, S. S. A. & Karim, Q. A. Omicron SARS-CoV-2 variant: a new chapter in the COVID-19 pandemic. Lancet, doi:10.1016/S0140-6736(21)02758-6 (2021).

41 Chen, R. T. E. et al. Resistance of SARS-CoV-2 variants to neutralization by monoclonal and serum-derived polyclonal antibodies. Nature Medicine 27, doi:10.1038/s41591-021-01294-w (2021).

42 Starr, T. N., Greaney, A. J., Dingens, A. S. & Bloom, J. D. Complete map of SARS-CoV-2 RBD mutations that escape the monoclonal antibody LY-CoV555 and its cocktail with LY-CoV016. Cell Rep Med 2, doi:ARTN 100255 10.1016/j.xcrm.2021.100255 (2021).

43 Starr, T. N. et al. SARS-CoV-2 RBD antibodies that maximize breadth and resistance to escape. Nature 597, 97-+, doi:10.1038/s41586-021-03807-6 (2021).

44 Tortorici, M. A. et al. Ultrapotent human antibodies protect against SARS-CoV-2 challenge via multiple mechanisms. Science 370, 950–957, doi:10.1126/science.abe3354 (2020).

45 Barnes, C. O. et al. SARS-CoV-2 neutralizing antibody structures inform therapeutic strategies. Nature 588, 682–687, doi:10.1038/s41586-020-2852-1 (2020).

46 Lei, C., Yang, J., Hu, J. & Sun, X. On the Calculation of TCID50 for Quantitation of Virus Infectivity. Virol Sin 36, 141–144, doi:10.1007/s12250-020-00230-5 (2021).

47 Burleson, F. G., Chambers, T. M. & Wiedbrauk, D. L. Virology: A Laboratory Manual. (Elsevier Science, 2014).

48 Reed, L. J. & Muench, H. A simple method of estimating fifty per cent endpoints. American journal of epidemiology 27, 493–497 (1938).

49 Wrapp, D. et al. Cryo-EM structure of the 2019-nCoV spike in the prefusion conformation. Science 367, 1260–1263, doi:10.1126/science.abb2507 (2020).

50 Otwinowski, Z. & Minor, W. Processing of X-ray diffraction data collected in oscillation mode. Methods Enzymol 276, 307–326 (1997).

51 McCoy, A. J. et al. Phaser crystallographic software. J Appl Crystallogr 40, 658–674, doi:10.1107/S0021889807021206 (2007).

52 Emsley, P. & Cowtan, K. Coot: model-building tools for molecular graphics. Acta Crystallogr D Biol Crystallogr 60, 2126–2132, doi:10.1107/S0907444904019158 (2004).

53 Adams, P. D. et al. PHENIX: a comprehensive Python-based system for macromolecular structure solution. Acta Crystallogr D Biol Crystallogr 66, 213–221, doi:10.1107/S0907444909052925 (2010).

54 Murshudov, G. N. et al. REFMAC5 for the refinement of macromolecular crystal structures. Acta Crystallogr D Biol Crystallogr 67, 355–367, doi:10.1107/S0907444911001314 (2011).

55 Collaborative Computational Project, N. The CCP4 suite: programs for protein crystallography. Acta Crystallogr D Biol Crystallogr 50, 760–763, doi:10.1107/S0907444994003112 (1994).

56 Pettersen, E. F. et al. UCSF Chimera--a visualization system for exploratory research and analysis. J Comput Chem 25, 1605–1612, doi:10.1002/jcc.20084 (2004).

57 Goddard, T. D. et al. UCSF ChimeraX: Meeting modern challenges in visualization and analysis. Protein Sci 27, 14–25, doi:10.1002/pro.3235 (2018).

58 Zheng, S. Q. et al. MotionCor2: anisotropic correction of beam-induced motion for improved cryo-electron microscopy. Nat Methods 14, 331–332, doi:10.1038/nmeth.4193 (2017).

59 Zhang, K. Gctf: Real-time CTF determination and correction. J Struct Biol 193, 1–12, doi:10.1016/j.jsb.2015.11.003 (2016).

60 Zivanov, J. et al. New tools for automated high-resolution cryo-EM structure determination in RELION-3. Elife 7, doi:10.7554/eLife.42166 (2018).

61 Walls, A. C. et al. Structure, Function, and Antigenicity of the SARS-CoV-2 Spike Glycoprotein. Cell 183, 1735, doi:10.1016/j.cell.2020.11.032 (2020).

62 Liu, L. H. et al. Potent neutralizing antibodies against multiple epitopes on SARS-CoV-2 spike. Nature 584, 450-+, doi:10.1038/s41586-020-2571-7 (2020).

63 van Heel, M. & Schatz, M. Fourier shell correlation threshold criteria. Journal of Structural Biology 151, 250–262, doi:10.1016/j.jsb.2005.05.009 (2005).

64 Scheres, S. H. W. & Chen, S. X. Prevention of overfitting in cryo-EM structure determination. Nature Methods 9, 853–854, doi:10.1038/nmeth.2115 (2012).

65 Wardemann, H. et al. Predominant autoantibody production by early human B cell precursors. Science 301, 1374–1377, doi:10.1126/science.1086907 (2003).

66 Kucukelbir, A., Sigworth, F. J. & Tagare, H. D. Quantifying the local resolution of cryo-EM density maps. Nat Methods 11, 63–65, doi:10.1038/nmeth.2727 (2014).

67 Saphire, E. O. et al. Crystal structure of a neutralizing human IGG against HIV-1: a template for vaccine design. Science 293, 1155–1159, doi:10.1126/science.1061692 (2001).

