## Supplementary figures and images for "Conformational Flexibility in Neutralization of SARS-CoV-2 by Naturally Elicited Anti-SARS-CoV-2 Antibodies"

### Supplemental Figure 1

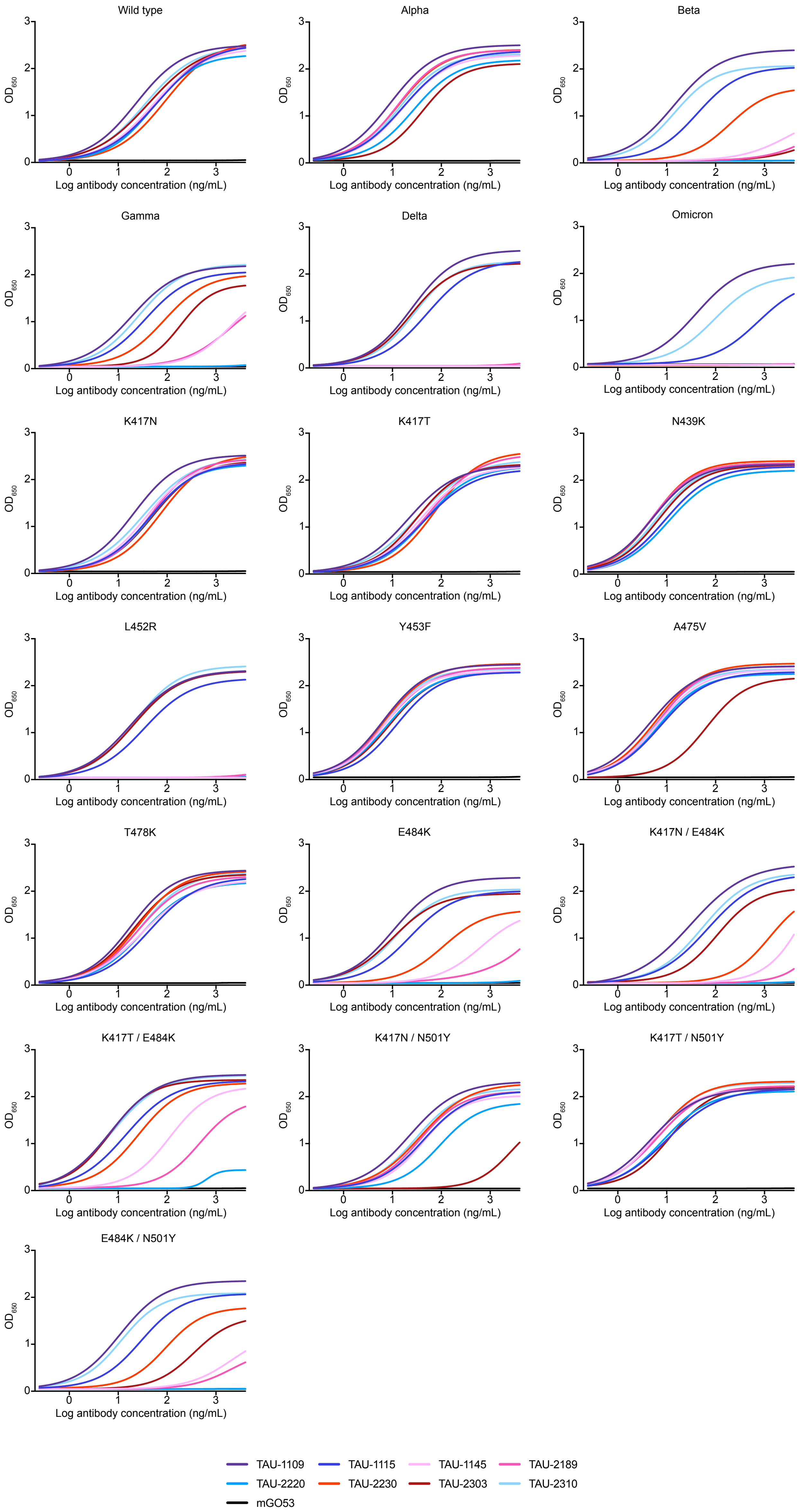

### Supplemental Figure 2

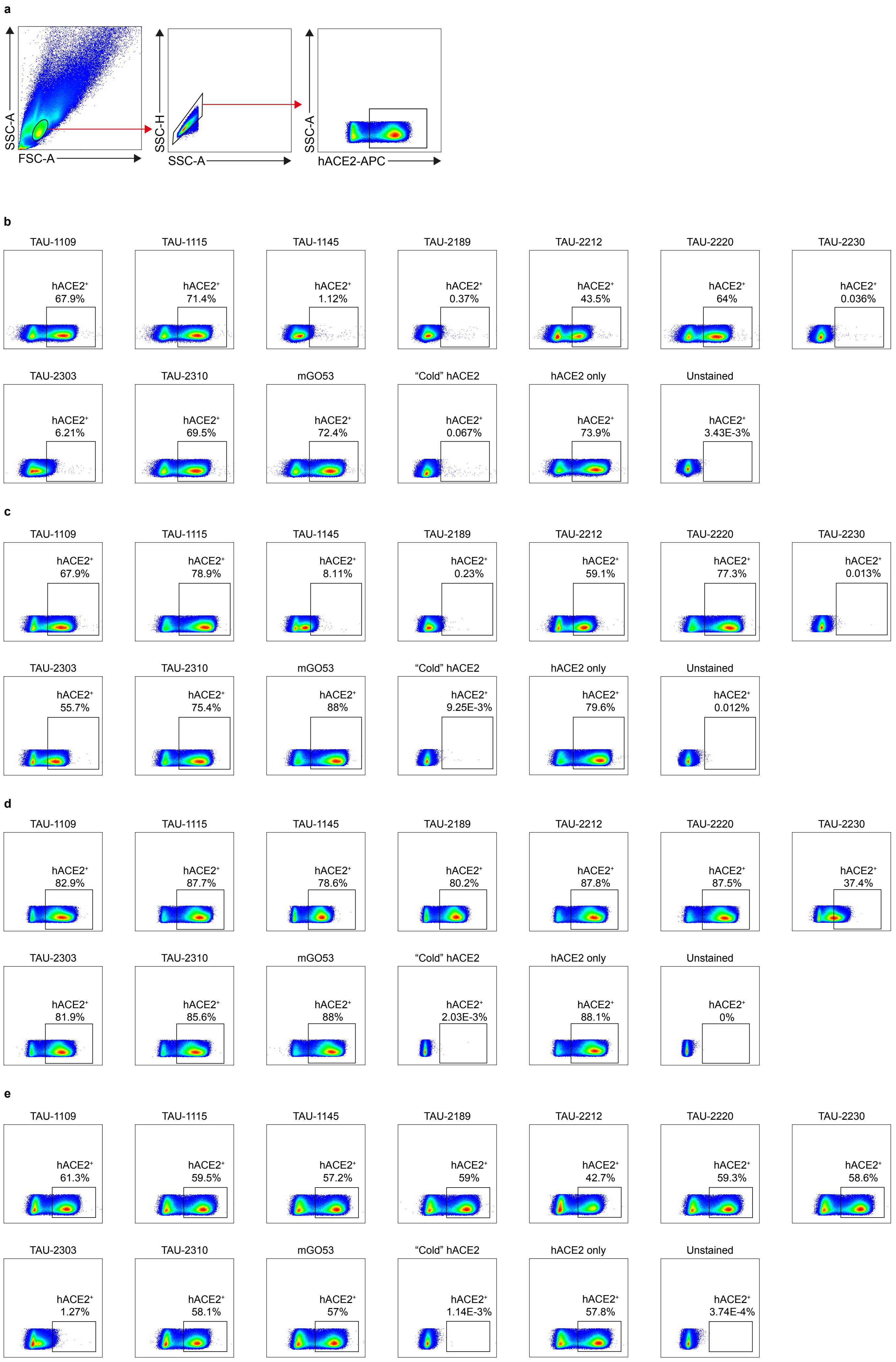

### Supplemental Figure 3

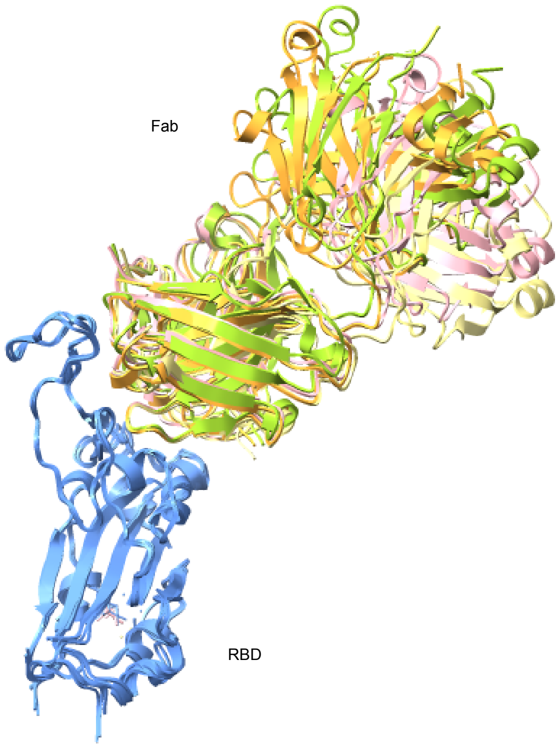

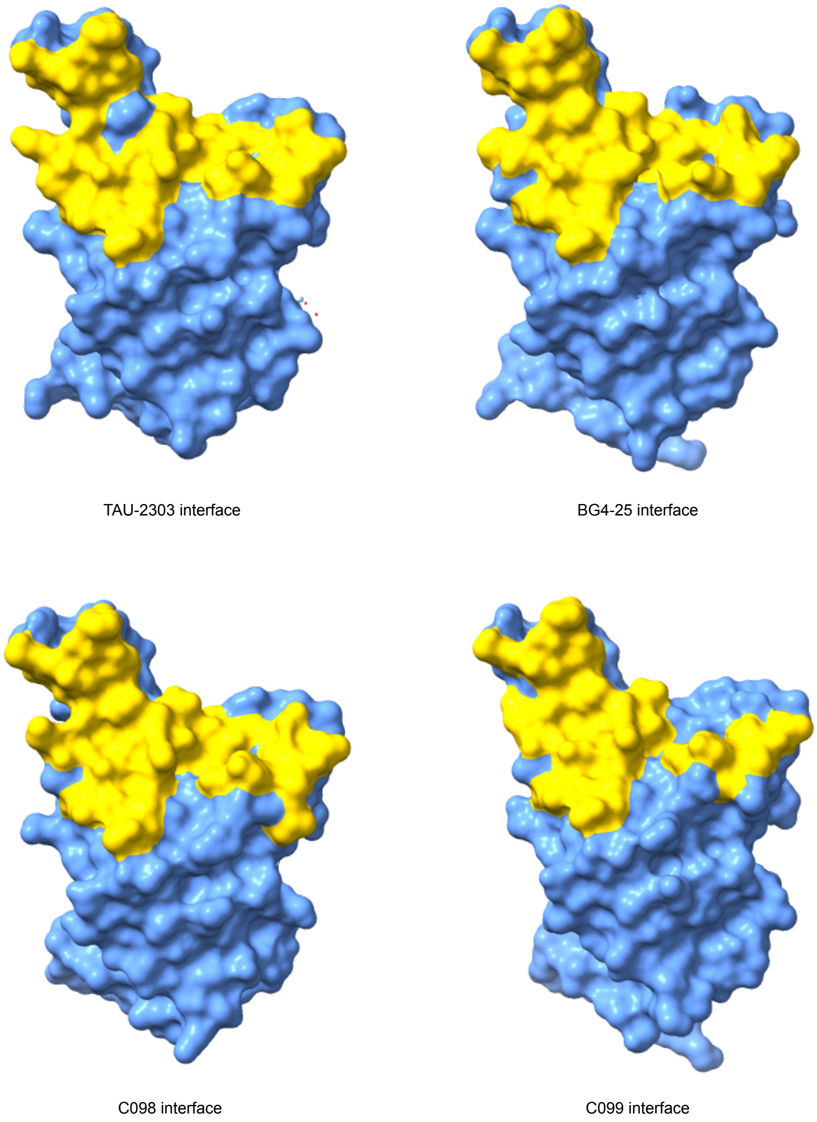

### Supplemental Figure 4

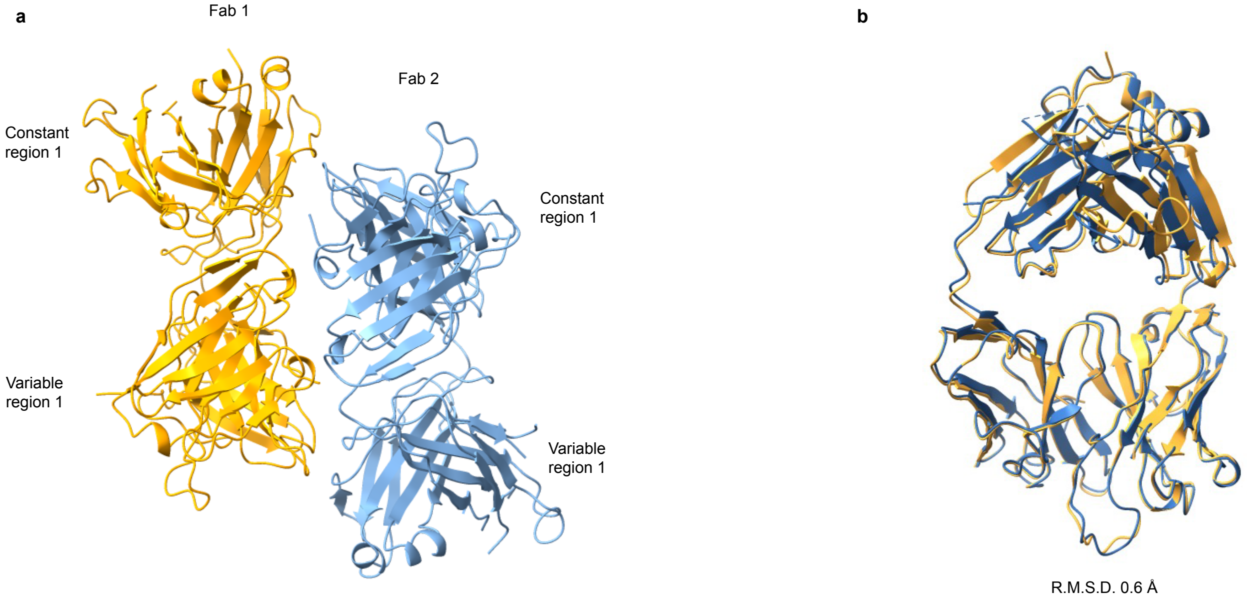

### Supplemental Figure 5

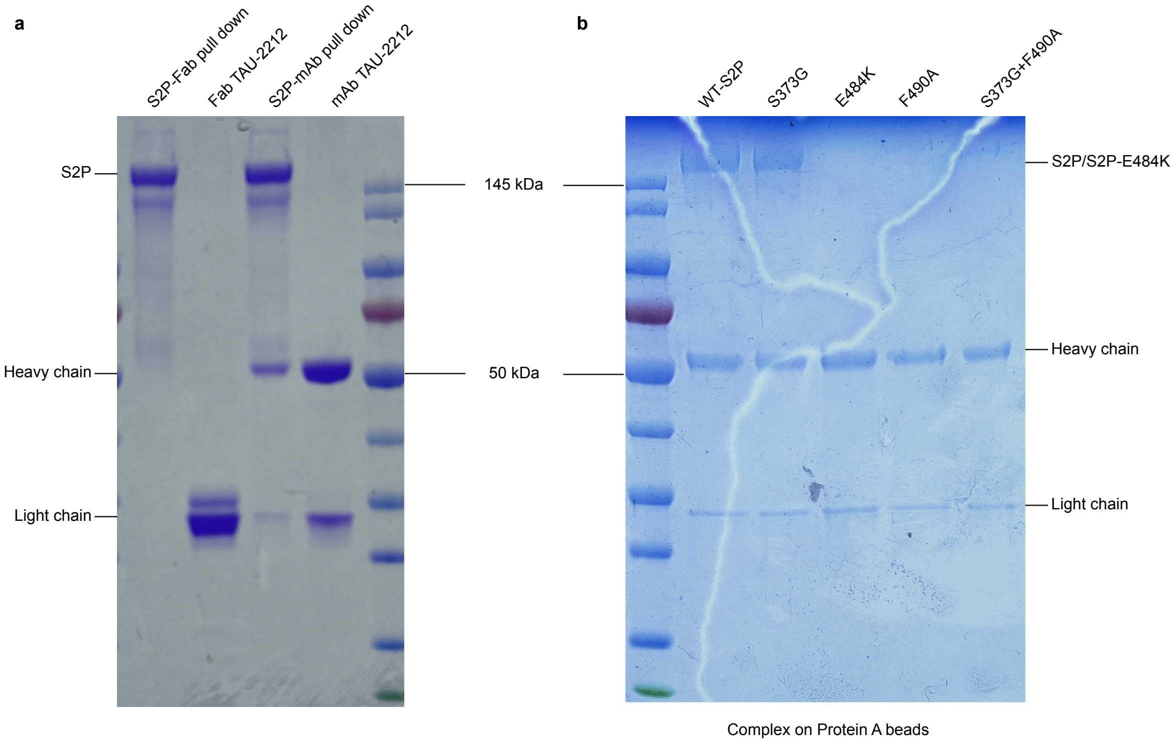

### Supplemental Figure 6

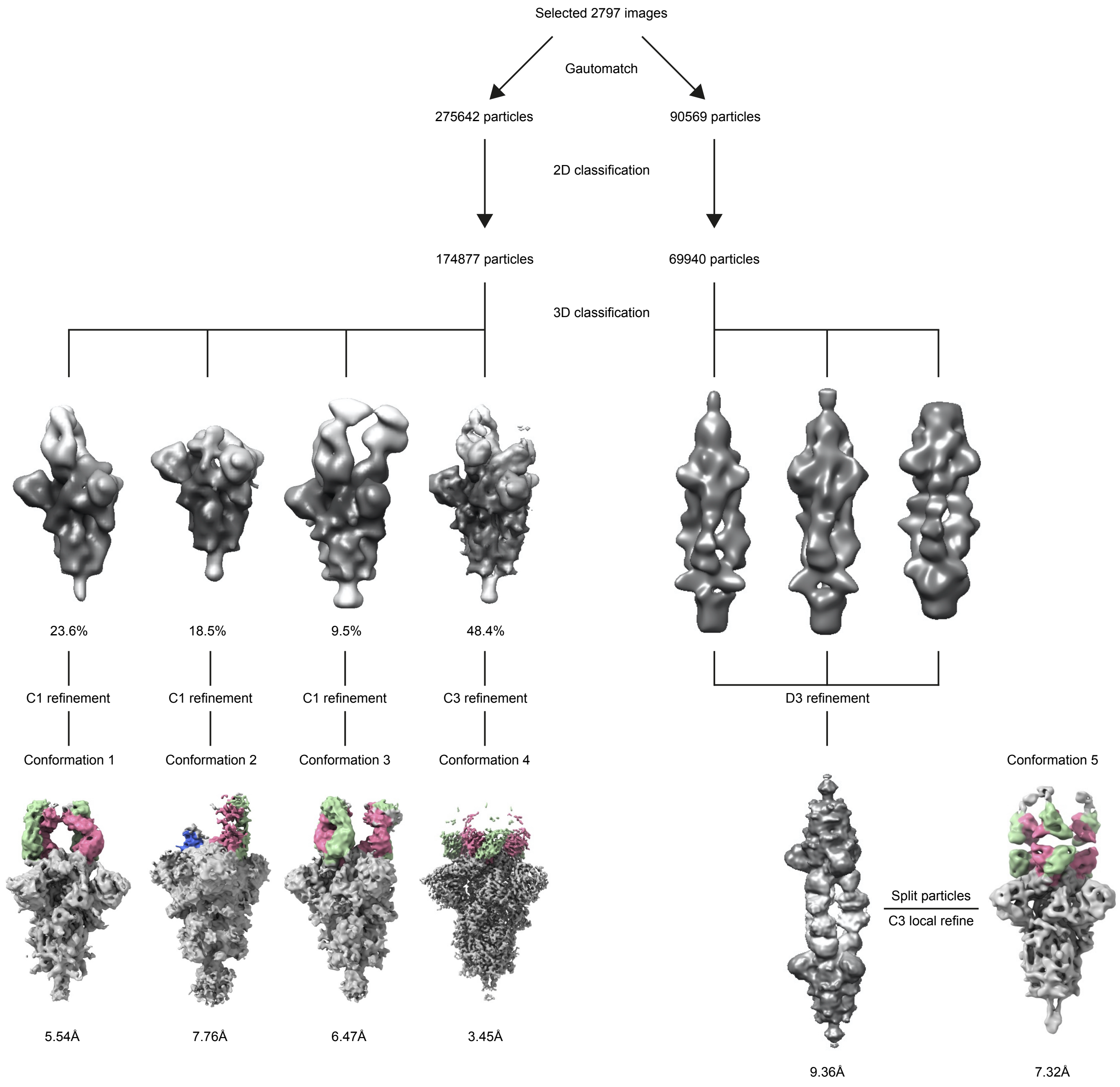

### Supplemental Figure 7

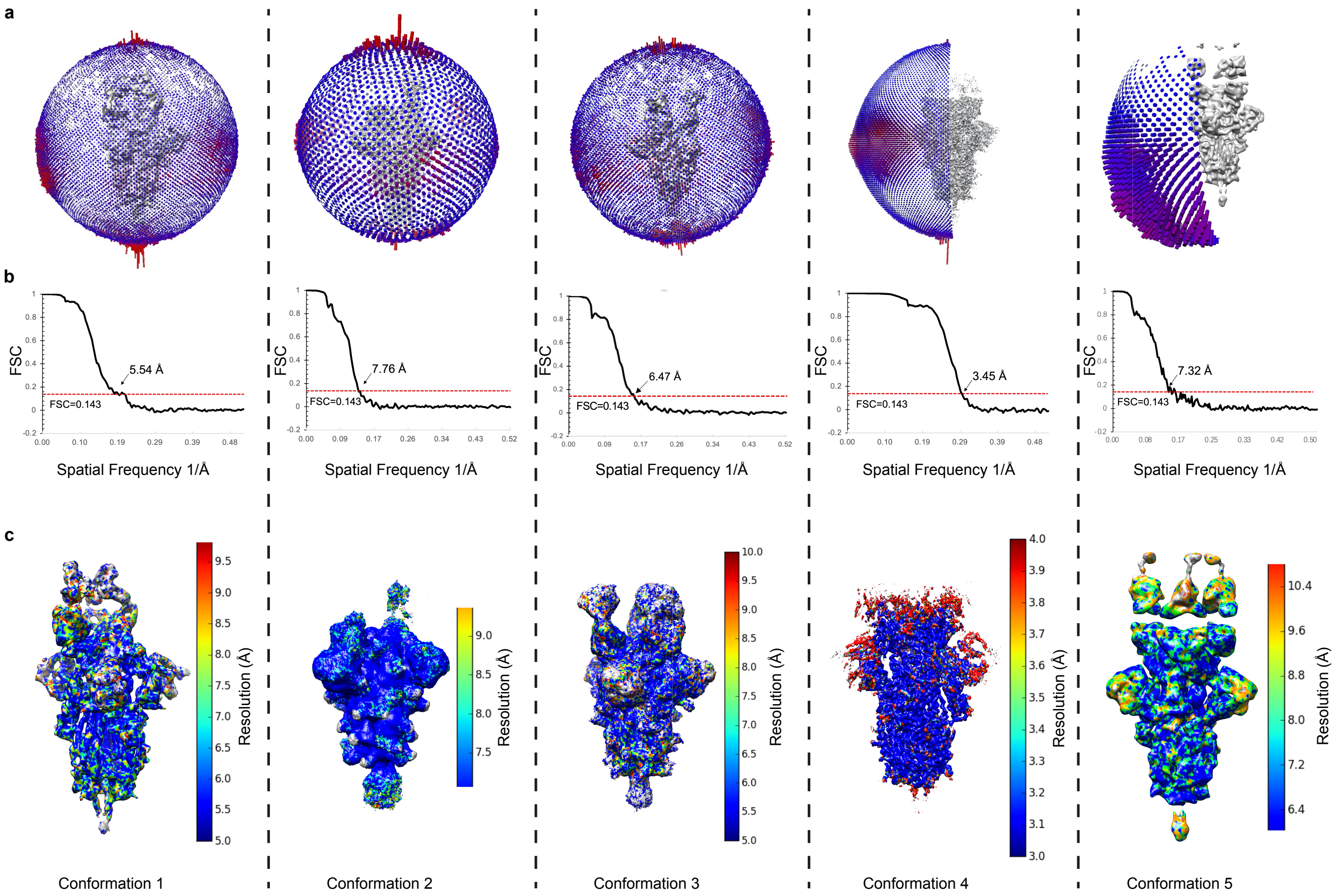

### Supplemental Figure 8

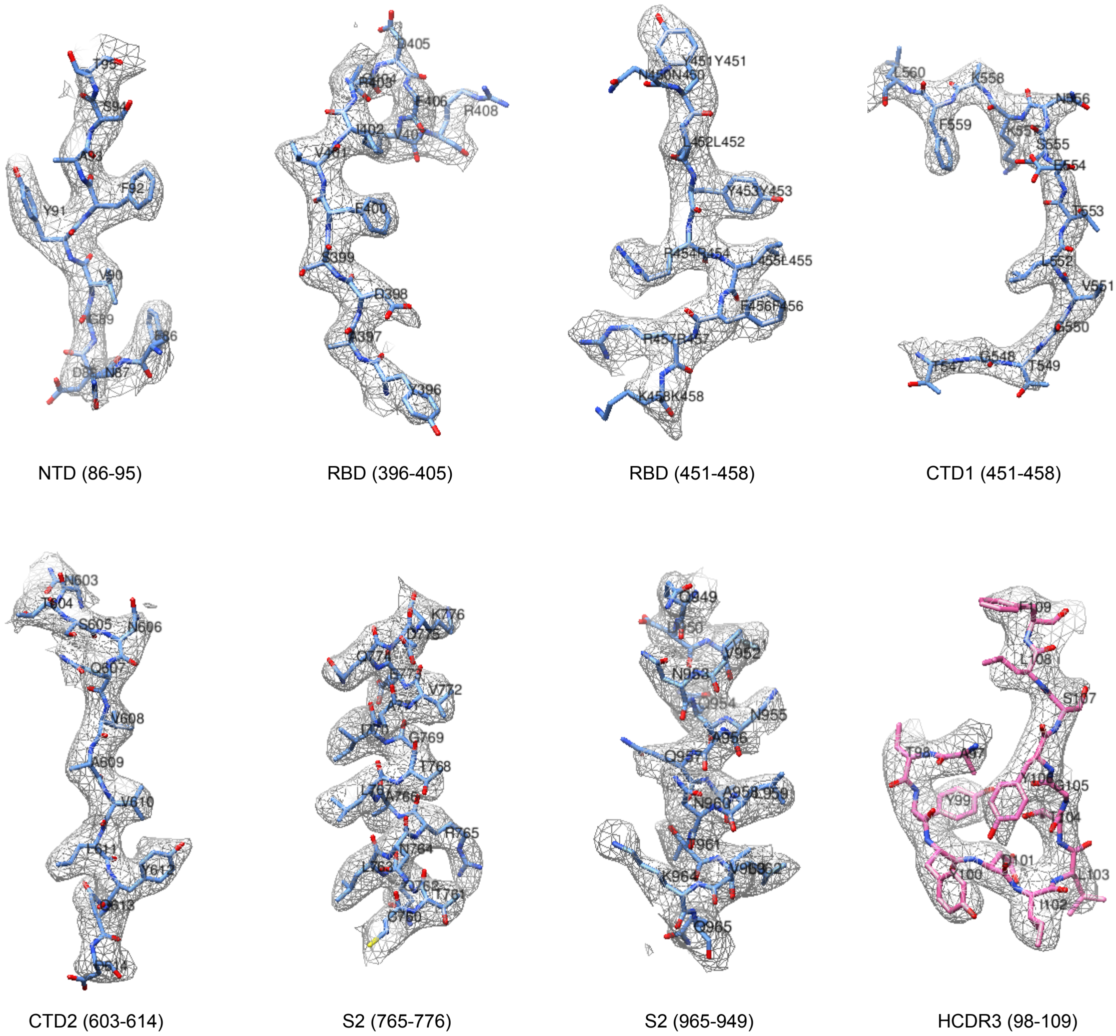

NTD (86-95) | RBD (396-405) | RBD (451-458) | CTD1 (451-458)

CTD2 (603-614) | S2 (765-776) | S2 (965-949) | HCDR3 (98-109)

### Supplemental Figure 9

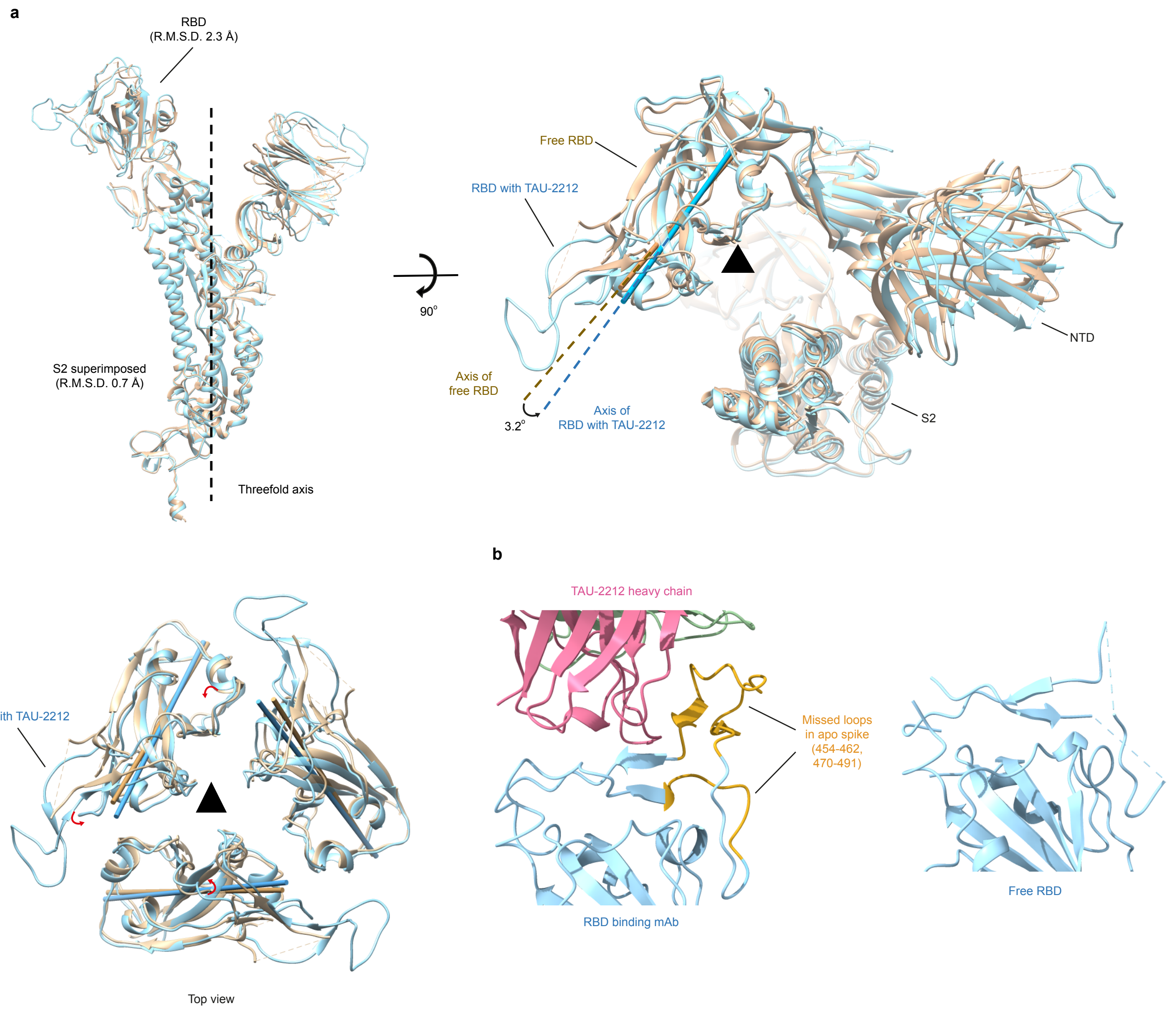

### Supplemental Figure 11

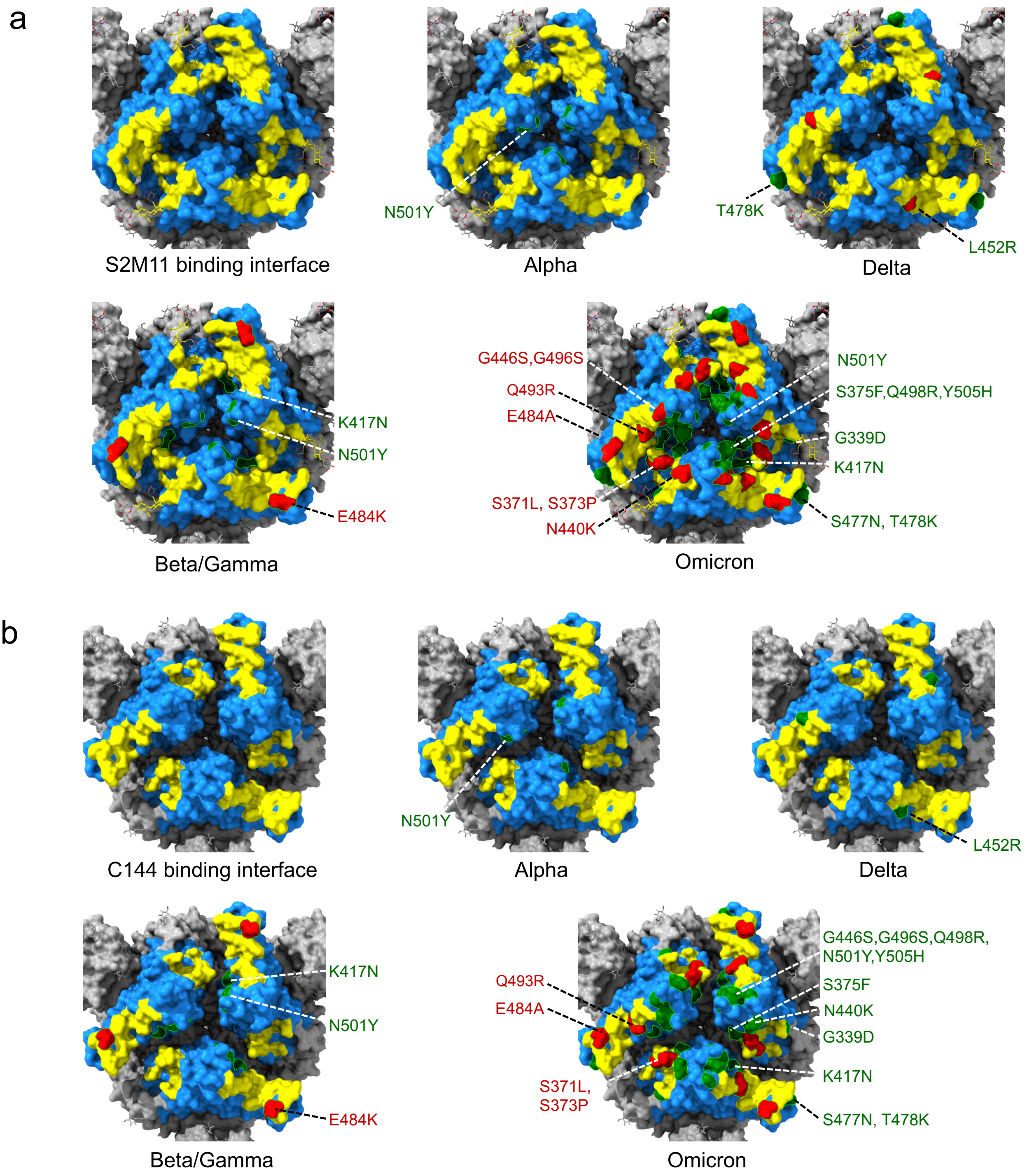
