## Supplemental Figure 10 for "Conformational Flexibility in Neutralization of SARS-CoV-2 by Naturally Elicited Anti-SARS-CoV-2 Antibodies"

**a**

```
                                    Fab
2212_H    QVQLVQSGAEVKKPGASVKVSCKASGYTFTGYYMHWVRQAPGQGLEWMGWINPNSGGTNY    60
1HZH_H    QVQLVQSGAEVKKPGASVKVSCQASGYRFSNFVIHWVRQAPGQRFEWMGWINPYNGNKEF    60
          **********************:**** *:.: :********** :******** .*..::

                                    Fab
2212_H    AQKFQGWVTMTRDTSISTAYMELSRLRSDDTAVYYCARGWATYY-DILTGYSLFDYWGQG    119
1HZH_H    SAKFQDRVTFTADTSANTAYMELRSLRSADTAVYYCARVGPYSWDDSPQDNYYMDVWGKG    120
          : ***. **:* *** .****** *** *********      : *    .   :* **:*

                                    Fab
2212_H    TLVTVSSASTKGPSVFPLAPSSKSTSGGTAALGCLVKDYFPEPVTVSWNSGALTSGVHTF    179
1HZH_H    TTVIVSSASTKGPSVFPLAPSSKSTSGGTAALGCLVKDYFPEPVTVSWNSGALTSGVHTF    180
          * * ********************************************************

                              Fab                          Loop
2212_H    PAVLQSSGLYSLSSVVTVPSSSLGTQTYICNVNHKPSNTKVDKRVEPKSCDKTHTCPPCP    239
1HZH_H    PAVLQSSGLYSLSSVVTVPSSSLGTQTYICNVNHKPSNTKVDKKAEPKSCDKTHTCPPCP    240
          *******************************************:.***************

                                    Fc
2212_H    APELLGGPSVFLFPPKPKDTLMISRTPEVTCVVVDVSHEDPEVKFNWYVDGVEVHNAKTK    299
1HZH_H    APELLGGPSVFLFPPKPKDTLMISRTPEVTCVVVDVSHEDPEVKFNWYVDGVEVHNAKTK    300
          ************************************************************

                                    Fc
2212_H    PREEQYNSTYRVVSVLTVLHQDWLNGKEYKCKVSNKALPAPIEKTISKAKGQPREPQVYT    359
1HZH_H    PREEQYNSTYRVVSVLTVLHQDWLNGKEYKCKVSNKALPAPIEKTISKAKGQPREPQVYT    360
          ************************************************************

                                    Fc
2212_H    LPPSREEMTKNQVSLTCLVKGFYPSDIAVEWESNGQPENNYKTTPPVLDSDGSFFLYSKL    419
1HZH_H    LPPSRDELTKNQVSLTCLVKGFYPSDIAVEWESNGQPENNYKTTPPVLDSDGSFFLYSKL    420
          *****:*:****************************************************

                          Fc
2212_H    TVDKSRWQQGNVFSCSVMHEALHNHYTQKSLSLSPGK    456
1HZH_H    TVDKSRWQQGNVFSCSVMHEALHNHYTQKSLSLSPGK    457
          *************************************
```

**b**

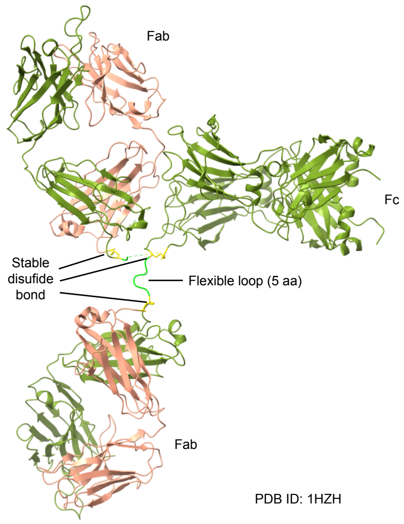


**c**

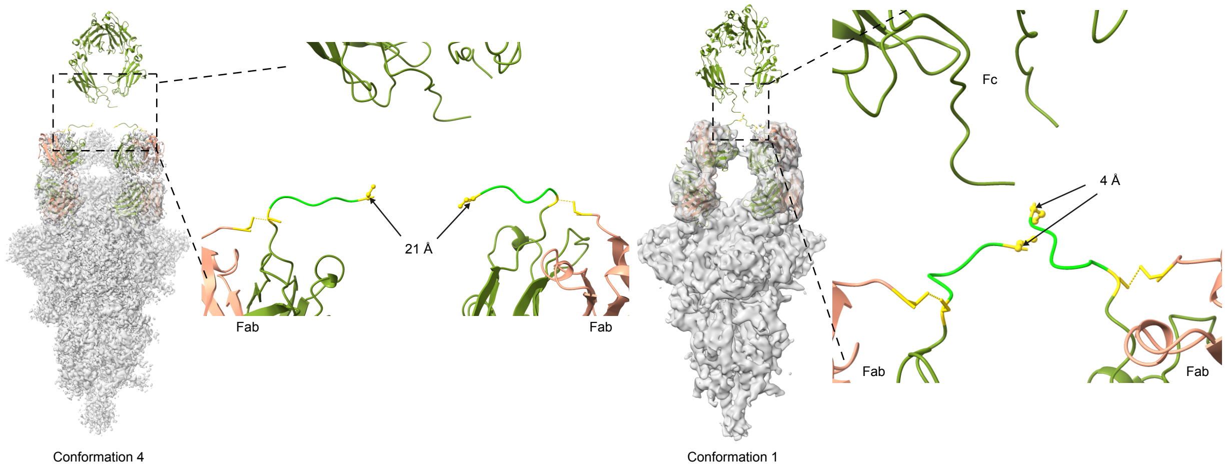


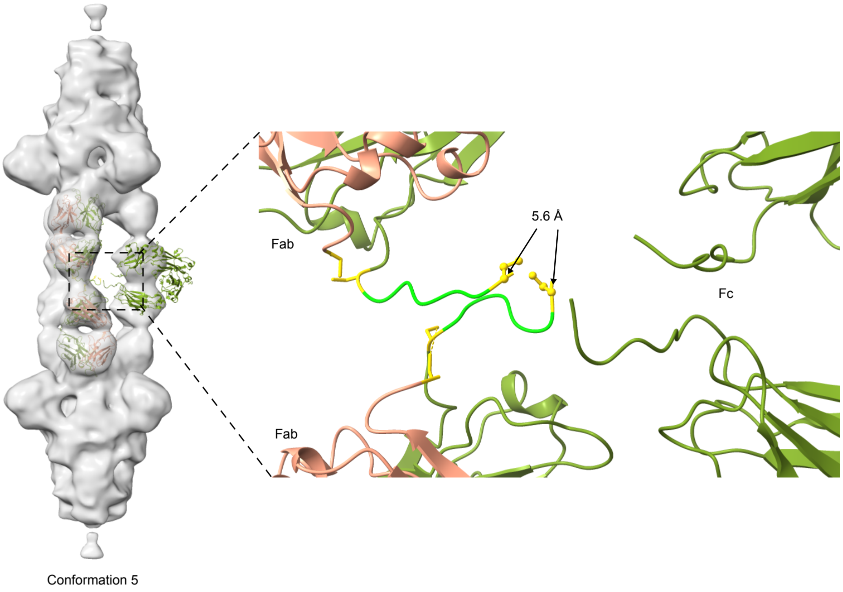
