## Supplemental Table 1 for "Conformational Flexibility in Neutralization of SARS-CoV-2 by Naturally Elicited Anti-SARS-CoV-2 Antibodies"

| | | Variant of concern | | | | |
|---|---|---|---|---|---|---|
| | Mutation | Alpha | Beta | Gamma | Delta | Omicron |
| | L18F | | | | | |
| | T20N | | | | | |
| | P26S | | | | | |
| | A67V | | | | | |
| | Δ69-70 | | | | | |
| | D80A | | | | | |
| | T95I | | | | | |
| | P138Y | | | | | |
| | G142D | | | | | |
| | Δ143 | | | | | |
| | Δ144-145 | | | | | |
| | R190S | | | | | |
| | N211I | | | | | |
| | Δ212 | | | | | |
| | 214EPEins | | | | | |
| | D215G | | | | | |
| | Δ242-244 | | | | | |
| | G339D | | | | | |
| | S371L | | | | | |
| | S373P | | | | | |
| | S375F | | | | | |
| | K417N | | | | | |
| | K417T | | | | | |
| | N440K | | | | | |
| | G446S | | | | | |
| RBD | L452R | | | | | |
| | S477N | | | | | |
| | T478K | | | | | |
| | E484A | | | | | |
| | E484K | | | | | |
| | Q493R | | | | | |
| | G498R | | | | | |
| | N501Y | | | | | |
| | Y505H | | | | | |
| | T547K | | | | | |
| | A570D | | | | | |
| | D614G | | | | | |
| | H655Y | | | | | |
| | N679K | | | | | |
| | P681H | | | | | |
| | P681R | | | | | |
| | A701V | | | | | |
| | T716I | | | | | |
| | N764K | | | | | |
| | D796Y | | | | | |
| | N856K | | | | | |
| | Q954H | | | | | |
| | N969K | | | | | |
| | L981F | | | | | |
| | S982A | | | | | |
| | T1027I | | | | | |
| | D1118H | | | | | |
| | V1176F | | | | | |
