## Supplemental Table 2 for "Conformational Flexibility in Neutralization of SARS-CoV-2 by Naturally Elicited Anti-SARS-CoV-2 Antibodies"

**Extended Data Table 2: X-ray data collection and refinement statistics for RBD-Fab2303 complex and Fab2212, Related to Figures 4 and 5.**

| | RBD-Fab2303 | Fab2212 |
|---|---|---|
| Data collection | | |
| Wavelength(Å) | 0.979 | 0.979 |
| Resolution(Å) | 20.09-2.42 (2.51-2.42) | 29.75-2.70 (2.80-2.70) |
| Space group | $C\ 2\ 2\ 2_1$ | $P\ 6_5$ |
| Unit cell | 85.13, 149.98, 144.79<br>90.0, 90.0, 90.0 | 75.977, 75.977, 348.14, 90, 90, 120 |
| Completeness (%) | 99.44 (95.98) | 99.67(97.56) |
| R-merge | 0.19 (0.751) | 0.145 (0.924) |
| Redundancy | 12.1 (10.2) | 8.5 (7.1) |
| I/σ (I) | 12 (2) | 25 (2.9) |
| Statistics for Refinement | | |
| No. of reflections | 35382 (3389) | 30759 (3004) |
| R-work | 0.1821 | 0.2464 |
| R-free | 0.2201 | 0.2716 |
| R.m.s.d | | |
| Bond(degree) | 0.72 | 0.97 |
| Length(Å) | 0.004 | 0.002 |
| Ramachandran plot | | |
| Favored region (%) | 97.24 | 94.99 |
| Allowed region (%) | 2.76 | 5.01 |
| Twin law | | k, h, -l |
| Twin fraction | | 0.41/0.59 |
