## Supplemental Table 3 for "Conformational Flexibility in Neutralization of SARS-CoV-2 by Naturally Elicited Anti-SARS-CoV-2 Antibodies"

### Extended Data Table 3: Contacts between RBD and Fab2303.

| Van der Waals contacts[a] | | Direct hydrogen bonds[b] | | |
|---|---|---|---|---|
| RBD | Fab 2303 | RBD | Fab 2303 | Distance (Å) |
| ARG$^{403}$ | ASN$^{L92}$(4), ASN$^{L93}$(1) | ARG$^{403}$[NH1] | ASN$^{L92}$[O] | 3.0 |
| THR$^{415}$ | SER$^{H56}$(2), TYR$^{H58}$(7) | THR$^{415}$[OG1] | TYR$^{H58}$[OH] | 2.7 |
| GLY$^{416}$ | TYR$^{H52}$(3), TYR$^{H58}$(2) | | | |
| ***LYS$^{417}$** | TYR$^{H33}$(2), TYR$^{H52}$(4), ASN$^{L92}$(1) | LYS$^{417}$[N] | TYR$^{H52}$[OH] | 3.6 |
| | | LYS$^{417}$[NZ] | TYR$^{H52}$[OH] | 3.6 |
| ASP$^{420}$ | TYR$^{H52}$(2), SER$^{H56}$(4) | ASP$^{420}$[OD2] | SER$^{H56}$[OG] | 2.5 |
| TYR$^{421}$ | TYR$^{H33}$(2), TYR$^{H52}$(5), SER$^{H53}$(4), GLY$^{H54}$(2) | TYR$^{421}$[OH] | SER$^{H53}$[N] | 3.3 |
| | | TYR$^{421}$[OH] | GLY$^{H54}$[N] | 3.1 |
| **TYR$^{453}$** | VAL$^{H101}$(1), ASN$^{L92}$(2) | TYR$^{453}$[OH] | ASN$^{L92}$[ND2] | 2.9 |
| **LEU$^{455}$** | TYR$^{H33}$(5), ALA$^{H100}$(2), VAL$^{H101}$(1) | LEU$^{455}$[O] | TYR$^{H33}$[OH] | 2.7 |
| ARG$^{457}$ | SER$^{H53}$(2) | | | |
| LYS$^{458}$ | ARG$^{H30}$(11), SER$^{H53}$(3), GLY$^{H54}$(2) | LYS$^{458}$[O] | ARG$^{H30}$[NH2] | 2.9 |
| | | SER$^{459}$[OG] | ARG$^{H30}$[NH1] | 3.3 |
| ASN$^{460}$ | GLY$^{H54}$(5) | | | |
| TYR$^{473}$ | SER$^{H31}$(6), SER$^{H53}$(2) | TYR$^{473}$[OH] | SER$^{H31}$[O] | 2.6 |
| | | TYR$^{473}$[OH] | SER$^{H53}$[OG] | 3.4 |
| GLN$^{474}$ | SER$^{H31}$(1) | | | |
| **ALA$^{475}$** | PHE$^{H27}$(2), THR$^{H28}$(2), ASN$^{H32}$(4) | ALA$^{475}$[O] | THR$^{H28}$[N] | 3.3 |
| | | ALA$^{475}$[O] | ASN$^{H32}$[ND2] | 3.0 |
| GLY$^{476}$ | THR$^{H28}$(2) | | | |
| SER$^{477}$ | THR$^{H28}$(1) | | | |
| **PHE$^{486}$** | VAL$^{H2}$(3), ARG$^{H97}$(2), ASP$^{H105}$(1) | | | |
| ***ASN$^{487}$** | GLY$^{H26}$(2), PHE$^{H27}$(2), ARG$^{H97}$(3) | ASN$^{487}$[ND2] | GLY$^{H26}$[O] | 2.9 |
| | | ASN$^{487}$[OD1] | ARG$^{H97}$[NH1] | 3.6 |
| | | ASN$^{487}$[OD1] | ARG$^{H97}$[NH2] | 3.2 |
| ***TYR$^{489}$** | ARG$^{H97}$(2), LEU$^{H99}$(5) | TYR$^{489}$[OH] | ARG$^{H97}$[NH1] | 3.6 |
| ***GLN$^{493}$** | VAL$^{H101}$(1), TYR$^{H102}$(4) | GLN$^{493}$[NE2] | TYR$^{H102}$[OH] | 2.7 |
| | | SER$^{494}$[O] | TYR$^{L32}$[OH] | 3.7 |
| TYR$^{495}$ | TYR$^{L32}$(5) | TYR$^{495}$[O] | TYR$^{L32}$[OH] | 2.9 |
| | | **GLY$^{496}$[O]** | SER$^{L30}$[OG] | 3.3 |

| | | | | |
|---|---|---|---|---|
| **GLN$^{498}$** | SER$^{L30}$(4),<br>SER$^{L67}$(3) | GLN$^{498}$[OE1] | SER$^{L30}$[OG] | 3.2 |
| | | GLN$^{498}$[OE1] | SER$^{L67}$[OG] | 3.3 |
| | | GLN$^{498}$[NE2] | SER$^{L67}$[OG] | 3.6 |
| ***THR$^{500}$** | GLY$^{L28}$(2) | | | |
| **ASN$^{501}$** | GLY$^{L28}$(4),<br>SER$^{L30}$(6) | ASN$^{501}$[OD1] | SER$^{L30}$[OG] | 2.8 |
| | | ASN$^{501}$[OD1] | SER$^{L30}$[N] | 3.4 |
| **GLY$^{502}$** | GLN$^{L27}$(2),<br>GLY$^{L28}$(5) | GLY$^{502}$[N] | GLY$^{L28}$[O] | 2.9 |
| ***TYR$^{505}$** | ILE$^{L2}$(2),<br>ILE$^{L29}$(1)<br>TYR$^{L32}$(4),<br>LEU$^{L91}$(1),<br>ASN$^{L92}$(2),<br>ASN$^{L93}$(2) | TYR$^{505}$[OH] | LEU$^{L91}$[O] | 2.9 |

a:Van der Waals contacts have interatomic distance ≤ 4.0 Å and defined by CCP4i.
b:Putative H-bonds and salt-bridges defined by PISA.
Residues in bold indicate the residues of RBD involved in the interactions with ACE2.
*Stars indicate RBD residues, of which the side chain atoms are involved in forming hydrogen bonds with ACE2.
