## Supplemental Table 4 for "Conformational Flexibility in Neutralization of SARS-CoV-2 by Naturally Elicited Anti-SARS-CoV-2 Antibodies"

**Extended Data Table 4: Cryo-EM data collection, refinement and validation statistics for RBD-Fab2303 and spike-TAU-2212.**

| | S6P - Fab 2303 | S2P - TAU-2212 | | | | |
|---|---|---|---|---|---|---|
| | | Conformation 1 | Conformation 2 | Conformation 3 | Conformation 4 | Conformation 5 |
| **Data collection and processing** | | | | | | |
| Voltage (kV) | 300 | 300 | | | | |
| Defocus range (μm) | -1.5 to -2.8 | -1.5 to -2.0 | | | | |
| Pixel size (Å) | 1.25 | 0.97 | | | | |
| Final particle images (no.) | 38331 | 20339 | 15868 | 14193 | 72056 | 39788 |
| Map resolution (Å) | 4.5 | 5.54 | 7.76 | 6.47 | 3.45 | 7.32 |
| Map sharpening B factor ($Å^2$) | -167.3 | -143.3 | -371.2 | -154.39 | -141.3 | -183.2 |
| Initial model used | 6VXX | 6VXX, 6XEY | | | | |
| Symmetry imposed | C1 | C1 | C1 | C1 | C3 | C3 |
| **Refinement** | | | | | | |
| R.M.S. deviations | | | | | | |
| Bond lengths (Å) | | | | | 0.003 | |
| Bond angles (°) | | | | | 0.547 | |
| MolProbity score | | | | | 1.76 | |
| Clashscore | | | | | 7.37 | |
| Rotamer outliers (%) | | | | | 0.17 | |
| Ramachandran plot | | | | | | |
| Favored (%) | | | | | 0.07 | |
| Allowed (%) | | | | | 5.12 | |
| Disallowed (%) | | | | | 94.81 | |
| B factors ($Å^2$) | | | | | | |
| Protein | | | | | 89.54 | |
| Ligand | | | | | 74.02 | |
