## Supplemental Table 5 for "Conformational Flexibility in Neutralization of SARS-CoV-2 by Naturally Elicited Anti-SARS-CoV-2 Antibodies"

**Extended Data Table 5: Contacts between spike and TAU-2212.**

| Van der Waals contacts[a] | | Direct hydrogen bonds[b] | | |
|---|---|---|---|---|
| Spike | Fab 2212 | Spike | Fab 2212 | Distance (Å) |
| VAL$^{445RBD1}$ | SER$^{H74}$(3) | | | |
| *ASN$^{448RBD1}$ | THR$^{H30}$(5),<br>THR$^{H73}$(1) | ASN$^{448RBD1}$[OD1] | THR$^{H30}$[OG1] | 2.9 |
| LEU$^{455RBD1}$ | TYR$^{H99}$(1) | | | |
| PHE$^{456RBD1}$ | TYR$^{H100}$(3) | | | |
| VAL$^{483RBD1}$ | PRO$^{L96}$(5) | | | |
| *GLU$^{484RBD1}$ | TYR$^{H33}$(5),<br>TRP$^{H50}$(3),<br>ASN$^{H52}$(1),<br>PRO$^{L96}$(2) | GLU$^{484RBD1}$[OE2] | TYR$^{H33}$[OH] | 2.5 |
| | | GLU$^{484RBD1}$[OE2] | ASN$^{H52}$[ND2] | 3.4 |
| GLY$^{485RBD1}$ | TYR$^{L90}$(2) | | | |
| *PHE$^{486RBD1}$ | TYR$^{H106}$(2),<br>TYR$^{L29}$(1),<br>LEU$^{L31}$(2),<br>TYR$^{L90}$(6) | PHE$^{486RBD1}$[N] | TYR$^{L90}$[OH] | 3.0 |
| TYR$^{489RBD1}$ | THR$^{H98}$(5),<br>TYR$^{H99}$(4),<br>TYR$^{H100}$(3) | | | |
| PHE$^{490RBD1}$ | ASN$^{H52}$(1),<br>SER$^{H54}$(4) | | | |
| *LEU$^{492RBD1}$ | ASN$^{H53}$(3) | LEU$^{492RBD1}$[O] | ASN$^{H53}$[ND2] | 3.5 |
| *GLN$^{493RBD1}$ | THR$^{H30}$(2),<br>ASN$^{H53}$(3),<br>THR$^{H98}$(1),<br>TYR$^{H99}$(1) | GLN$^{493RBD1}$[NE2] | THR$^{H30}$[O] | 3.6 |
| | | GLN$^{493RBD1}$[OE1] | THR$^{H98}$[OG1] | 3.3 |
| | | GLN$^{493RBD1}$[OE1] | ASN$^{H53}$[ND2] | 3.1 |
| *SER$^{494RBD1}$ | THR$^{H30}$(4),<br>ASN$^{H53}$(2) | SER$^{494RBD1}$[N] | ASN$^{H53}$[OD1] | 3.8 |
| | | SER$^{494RBD1}$[OG] | ASN$^{H53}$[OD1] | 3.3 |
| PHE$^{342RBD2}$ | LEU$^{H103}$(1) | | | |
| VAL$^{367RBD2}$ | TYR$^{H100}$(3) | | | |
| ASN$^{370RBD2}$ | TYR$^{H100}$(3) | | | |
| *SER$^{371RBD2}$ | TYR$^{H100}$(2),<br>ASP$^{H101}$(3),<br>ILE$^{H102}$(1) | SER$^{371RBD2}$[OG] | ILE$^{H102}$[N] | 3.7 |
| *ALA$^{372RBD2}$ | TYR$^{H99}$(1),<br>TYR$^{H100}$(2) | ALA$^{372RBD2}$[N] | TYR$^{H100}$[O] | 3.2 |
| *SER$^{373RBD2}$ | ASP$^{H101}$(7) | SER$^{373RBD2}$[OG] | ASP$^{H101}$[OD1] | 3.1 |
| | | SER$^{373RBD2}$[N] | ASP$^{H101}$[2D1] | 3.8 |
| PHE$^{374RBD2}$ | ILE$^{H102}$(1),<br>LEU$^{H103}$(1) | | | |

a:Van der Waals contacts have interatomic distance ≤ 4.0 Å and defined by CCP4i.
b:Putative H-bonds and salt-bridges defined by PISA.
In the interface, RBD1 binds both with heavy and light chains. RBD2 only binds heavy chain.
H, heavy chain of mAb TAU-2212. L, light chain of mAb TAU-2212
*Stars indicate RBD residues, of which the side chain atoms are involved in forming hydrogen bonds with ACE2.
